# Cyclin-dependent kinase 9 inhibitors as oncogene signaling modulators in combination with targeted therapy for the treatment of colorectal cancer

**DOI:** 10.1101/2025.08.13.669992

**Authors:** Mahshid Mohammadi, Muzaffer Ahmed Bhat, Terence Li, Ning Wei, Priyanka Patil, Christiana Mo, Natalie Thielen, Zimo Huang, Juan Du, Doctor Yitzchak Goldstein, Othon Wiltz, Renee Huang, Kohtaro Ooka, Melanie Quintal, Edward Chu, Chaoyuan Kuang

**Affiliations:** Montefiore Einstein Comprehensive Cancer Center, Bronx, NY; Department of Oncology, Montefiore Einstein College of Medicine, Bronx, NY; Department of Molecular Pharmacology, Montefiore Einstein College of Medicine, Bronx, NY; Marilyn and Stanley M. Katz Institute for Immunotherapy for Cancer and Inflammatory Disorders, Bronx, NY; Department of Pathology, Montefiore Einstein College of Medicine, Bronx, NY; Department of Cell Biology, Montefiore Einstein College of Medicine, Bronx, NY; Department of Surgery, Montefiore Einstein College of Medicine, Bronx, NY; Division of Gastroenterology, Department of Medicine, Montefiore Einstein College of Medicine, Bronx, NY

**Keywords:** Colorectal cancer, drug development, CDK9 inhibitors, targeted therapy, patient-derived organoids

## Abstract

**Background:** Colorectal cancer (CRC) is the second deadliest cancer worldwide and new treatment options are urgently needed. Cyclin dependent kinase 9 (CDK9) promotes aberrant RNA transcription in cancer and is a promising target for cancer therapies.

**Methods:** Using CRC cell lines as well as newly established patient-derived organoid models of CRC, we studied the clinically promising CDK9 inhibitors (AZD4573, BAY1125152/VIP152/enitociclib, and NVP2) to determine their therapeutic potential. We investigated the efficacy and mechanisms of action through cell growth and cytotoxicity assays, RNAseq, immunoblotting, and IHC.

**Results:** We found CDK9 inhibitors to be highly potent against CRC, through suppression of proliferation and induction of apoptosis. Our results demonstrated CDK9 inhibitors to be broadly active against a set of CRC models derived from a diverse patient population. Our mechanistic studies showed significant suppression of the Mitogen-active protein kinase (MAPK) signaling pathway due to CDK9 inhibitor treatment, suggesting that CDK9 inhibitor efficacy could be enhanced when combined with MAPK pathway inhibitors. As proof-of-concept, we found that CDK9 inhibitors and MEK inhibitors could be combined to synergistically suppress CRC growth and survival.

**Conclusions:** CDK9 inhibitors show promising activity against patient-derived models of CRC. MAPK signaling is particularly suppressed by CDK9 inhibitors. Combining CDK9 inhibitors and targeted therapy against MAPK signaling pathway may be a viable strategy worthy of further investigation preclinically and clinically.

**State of translational relevance:** Novel therapies for chemotherapy-refractory CRC remain urgently needed. Rapid validation of efficacy in appropriate translational models can provide the necessary rationale to test new therapies in clinical trials. Using newly established patient-derived organoids, we validate the preclinical efficacy of transcriptional CDK9 inhibitors against CRC. This is a therapeutic class that has been scarcely evaluated for efficacy against CRC. We found that CDK9 inhibitors potently inhibit patient-derived organoid models of CRC. Our mechanistic work also reveals that clinically achievable concentrations of CDK9 inhibitor treatment results in suppression of MAPK signaling. As proof-of-concept of the potential actionability of our findings, we demonstrate that CDK9 inhibitors can be combined with MEK inhibitors to synergistically suppress CRC growth. Our study provides new patient-derived CRC models as a resource available to others for therapeutic testing, as well as a roadmap for novel hypotheses about synergistic combination treatments using CDK9 inhibitors.

## INTRODUCTION

Colorectal cancer (CRC) is the third most common cancer and second deadliest cancer worldwide^1,2^. Combination chemotherapy and targeted therapies are frequently used to treat metastatic CRC (mCRC)^3,4^. First- and second-line therapy includes well-established cytotoxic drugs such as 5-fluorouracil or capecitabine, combined with oxaliplatin or irinotecan. These chemotherapy doublets are often combined with the anti-EGFR antibodies panitumumab or cetuximab, or the anti-VEGF antibody bevacizumab. Third-line treatment and beyond typically consists of oral agents such as regorafenib, trifluridine/tiparacil plus bevacizumab, or fruquintinib. In rare cases of microsatellite instability high (MSI-H) mCRC, treatment with immune checkpoint inhibitors can lead to significant durable responses^5,6^. However, most patients are microsatellite stable (MSS) where immunotherapy has only limited activity, and MSS patients need to be treated with cytotoxic chemotherapy as the backbone of their therapy. These patients usually progress through multiple standard-of-care chemotherapy regimens, and they eventually succumb to their disease. As a result, novel therapies for mCRC are urgently needed.

Genomic instability and epigenetic reprogramming are among the well-established Hallmarks of Cancer^7,8^. As a result of these characteristics, cancers including CRC have significantly altered RNA transcriptional landscapes as compared to non-malignant normal tissue. Dysfunctional gene expression then leads to a plethora of additional cellular consequences, which overlap with other Hallmarks of Cancer^9,10^. The importance of transcriptional dysregulation is further highlighted by the categorization of CRC into distinct consensus molecular subtypes (CMS) based on their transcriptional profiles. Each CMS group has been correlated with unique biological and clinical features, such as increased anti-tumor immune response (CMS1), increased chromosomal instability (CMS2), or increased TGF-β signaling (CMS4)^11^. Given the importance of transcriptional dysregulation in CRC biology, we reasoned that inhibition of pathogenic transcription may be a viable therapeutic strategy for CRC.

Cyclin-dependent kinase 9 (CDK9) is a protein kinase whose canonical function is to facilitate the transition of RNA Polymerase II (RNA Pol II) from the poised, promoter-paused state, into the elongation state. CDK9 phosphorylates serine residues on the C-terminal domain of RNA Pol II to achieve the transition to RNA elongation^12^. Numerous oncogenes have been shown to be driven by aberrant transcription, in part due to increased CDK9 activity^13–16^. Most notably, Mixed lineage leukemia 1 (MCL-1) and c-MYC are dependent on CDK9-driven transcriptional upregulation, especially in hematologic malignancies^15,17^. Given the importance of transcriptional upregulation, and specifically CDK9 activity, in driving carcinogenesis, we investigated the potential role of CDK9 as a therapeutic target for CRC.

## MATERIALS AND METHODS

Please see key resource tables (Supplementary Tables 1,2) for detailed information about key reagents.

### Cell lines

HCT-116 (RRID:CVCL_0291), DLD-1 (RRID:CVCL_0248), L-WRN (RRID:CVCL_DA06), and RKO (RRID:CVCL_0504) were purchased from ATCC. Lim1215 (RRID:CVCL_2574) was purchased from Sigma-Aldrich. HCT-116 was cultured in McCoy’s 5A (Gibco, 16600108) media supplemented with 10% FBS and 1% penicillin-streptomycin. DLD-1, RKO, and Lim1215 were cultured in RPMI1640 (Corning, MT10040CV) supplemented with 10% FBS and 1% penicillin-streptomycin, and L-WRN were cultured in DMEM supplemented with 10% FBS and 1% penicillin-streptomycin. All cell lines were maintained at 37 °C with 5% CO_2_ and grown with 100 U/mL penicillin and 100 μg/mL streptomycin. All experiments were conducted in cell line passages 2-15. All cell lines were monitored for *Mycoplasma* infection and treated as needed. HLA/STR testing was used to validate cell lines.

### Patient specimens

All patients were consented and registered on the IRB-approved Montefiore Einstein Comprehensive Cancer Center Biobank protocol (Einstein IRB# 2021-13730) prior to tissue and data acquisition. Clinical recruitment, coordination, and regulatory assistance were provided by the Cancer Clinical Trials Office (CCTO) of the Montefiore Einstein Comprehensive Cancer Center (MECCC). Tissues were obtained from patients who underwent standard-of-care surgical resection procedures, after the resection was completed and gross tumor specimen had been examined by pathology lab staff. Specimens were inspected, divided, and portions were used to immediately (1) fix in 10% neutral buffered formalin, (2) used to establish patient-derived organoids, and (3) frozen in DMEM supplemented with 20% FBS and 10% DMSO.

### Patient-derived organoids

Biobank participants were identified by treating physicians or surgeons, consented and enrolled in the biobank, and fresh tissue from surgical resections of CRC were obtained for investigation. Approximately 0.5 cm fragments of tumor tissue were used to establish PDOs. In brief, tissue fragments were mechanically and chemically disrupted using a razor blade and diluted gentle Collagenase/Hyaluronidase (Stemcell Technology, #07919), respectively. Tumor clusters were embedded in Matrigel and surrounded by complete PDO media, which was supplemented with WRN conditional media as well as ROCK inhibitor (Y-27632, Cayman Chemical, #10005583). All PDOs were passaged at least twice and growing with a doubling time of 3-10 days. For full description of PDO culture methods, see Supplementary Methods.

### Drug treatment

All drug treatments were performed by dilution of DMSO drug stock into aqueous media, with final DMSO concentrations at 0.1% or lower. Drug-free controls were used with a control DMSO concentration of 0.1%. Drug-free growth media was removed and replaced with drug-containing or DMSO-containing media at indicated concentrations. For PDO treatment, Matrigel+PDO domes were left intact while drug-free media was exchanged for drug-media, as described above.

### Cell viability assays

As previously described^18^, cells were plated in 96-well plates, allowed 24 h for attachment, and then treated with drugs for 48-72 hours. Cells were then assayed using a chemical viability assay (MTS, Promega G3581). PDO viability: PDOs were grown to confluency, then digested and filtered through a 40 µm filter to eliminate large tumor clusters and debris. PDOs were then resuspended in fresh Matrigel and plated in 10 µL aliquots per well in a 96-well plate. PDOs were allowed to grow and re-establish for 3-5 days, followed by drug treatment for 5 days.

Viability was measured using CellTiter-Glo 3D assay (Promega G9683). All viability assays were measured using the FLUOstar Omega microplate reader (BMG Labtech). All assays were performed with technical triplicate and repeated three times for biological triplicates. Synergy scores were calculated using the Chou-Talalay method^19^.

### Colony formation assay

As previously described^18^, colony formation was assayed by plating equal numbers of 24-hour drug-treated cells in 6-well plates at appropriate dilutions with no drug, followed by visualization and quantification of colonies by crystal violet staining after 14 days.

### Apoptosis assays

Cells were cultured and treated for 48 hours with inhibitors. Apoptosis was evaluated with FITC Annexin V Apoptosis Detection Kit (BD Biosciences #556570) and PI. The apoptotic cells were quantified with Invitrogen Attune NxT flow cytometry and FlowJo software (Version 10.1, RRID:SCR_008520) and calculated as percentages using GraphPad Prism version 9.4.1 software (RRID:SCR_002798).

### Cell cycle profile

HCT116 cells were plated at 30%-40% density and treated with inhibitors for 4 or 8 h. DMSO at 0.1% was used as vehicle control. The cells were then fixed with ice cold 70% ethanol for 4 h at 4 °C. RNase A (100 µg/mL) (Thermo Scientific #EN0531) and PI (0.05 µg/mL, BioLegend #421301) were added to cells. The stained cells were detected by Invitrogen NxT Attune flow cytometer and analyzed using FlowJo software (version 10.1) and calculated as percentage using GraphPad Prism version 9.4.1 software. Cell cycle profile curves were optimized and fitted to root-mean-squared differentials (FlowJo Version 10.1).

### Immunoblotting

Cells/PDOs were digested using 100 µL of RIPA (Boston Bioproducts NC9193720) supplemented with Protease Inhibitor Cocktail (Sigma P8340). Prior to digestion, PDOs were treated with cell recovery solution (Fisher 8774405) to remove Matrigel. Cells were incubated on ice for 10 minutes, then sonicated (Fisher, 422-A) for 5 seconds on and 5 seconds off for a total 15 seconds on at 20% AMP. Cleared lysates were mixed with 4X Laemmli (BioRad, 1610747) with 2-Mercaptoethanol (Sigma, M7522) and heated at 85°C for 10 minutes. The final lysates were then loaded into 4–15% Mini-PROTEAN® TGX™ gels (BioRad 4561086) alongside Precision Plus Protein™ Dual Color Standards ladder (BioRad 1610374). Gels were run with 1X running buffer (BioRad 1610772) at 160V for 35-45 minutes. Protein was transferred from gels onto activated PVDF membrane (Millipore IPVH00010) using a TransBlot Turbo (BioRad 1704150) and 1X transfer buffer (BioRad 1610771), methanol, and 10% sodium dodecyl sulfate (Lonza 51213).

Membranes were blocked for 1 hour in 1X TBS (BioRad 1706435) and Tween20 (TBST) supplemented with 10% milk (BioRad 1706404). Blocked membranes were incubated in primary antibody diluted in TBST with 10% milk, overnight while rocking in 4°C. Membranes were washed 3 times with TBST for 6 minutes each wash, then incubated with HRP-conjugated secondary antibody in TBST + 10% milk for 1 hour at room temperature. ECL solution (BioRad 1705061) was mixed fresh and added to the membranes, followed by imaging in a Li-Cor Odyssey FC (Li-Cor Biosciences). Densitometry of protein bands were measured using ImageJ (version 1.54G, RRID:SCR_003070) and normalized to GAPDH housekeeping protein from the same blot/membrane. See Supplementary Table 2 for primary antibodies and conditions.

### Reverse Transcriptase Quantitative Real-Time PCR (RT-qPCR)

Total RNA was isolated from patient tissue specimens, cell lines, and PDOs using the Direct-zol RNA MiniPrep RNA Isolation Kit (Zymo Research, R2071) according to the manufacturer’s protocol. RNA was reverse-transcribed into cDNA using the iScript™ Advanced cDNA Synthesis Kit (Bio-Rad, 1725037) according to the manufacturer’s instructions (Bio-Rad T100 cycler).

SsoAdvanced™ Universal SYBR Green Supermix (Bio-Rad, 1725270) was used according to the manufacturer’s protocol for pPCR amplification. Reactions were carried out on a CFX Opus 384 Real-Time PCR System (Bio-Rad, RRID:SCR_017251). The primers used for each gene are listed in Supplementary Table 1.

### Light microscopy

All brightfield and phase contrast micro-photographs were obtained using the Echo Revolve 4 combination inverted/upright microscope (Echo).

### RNA sequencing

For RNA sequencing (RNAseq) of cell lines, primary tumors, and PDOs, total RNA was extracted using the Direct-zol RNA MiniPrep Kit (Zymo Research, Cat # R2071) following the manufacturer’s protocol. Sequencing was performed on the NovaSeq 6000 platform (PE150) using poly-A capture and non-directional library preparation. Differential expression analysis between two or more sample groups was conducted using DESeq2 (https://www.bioconductor.org/packages, RRID:SCR_000154). Gene Ontology (GO) analysis and gene set enrichment analysis (GSEA) of the DEGs were performed using Parametric Analysis of Gene Set Enrichment (PAGE) implemented in the PGSEA package (https://www.bioconductor.org/packages). Enrichment significance was determined using Fisher’s exact test. For full RNAseq methods, see Supplementary Methods.

### Genomic sequencing

Genomic DNA was extracted from five primary tumor samples and six corresponding PDOs (passages 5–10, one replicate each) using the Quick-DNA/RNA™ MiniPrep Plus Kit (Zymo Research) following the manufacturer’s protocol. Whole exome sequencing (WES) was performed on Illumina platforms using a paired-end 150 bp (PE150) strategy. Reads were aligned to the human reference genome (hg19) using the Burrows-Wheeler Aligner (BWA, v0.7.17, RRID:SCR_010910). Somatic SNP, InDel, and CNV calling were performed using the Genome Analysis Toolkit (GATK, v4.3.0, RRID:SCR_001876), Mutect2 (RRID:SCR_026692), and Control-FREEC (v11.4, RRID:SCR_010822). Annotation of variants was conducted with ANNOVAR (RRID:SCR_012821). A complementary approach for tumor genomic characterization was also used for tumor tissue and PDO genomic sequencing. In brief, either matched adjacent normal tissue or Agilent OneSeq Human Reference DNA (male: 5190-8848, female: 5190-8850) was used alongside tumor and PDO DNA. Library preparation was performed using the SureSelectXT platform (Agilent Technologies), and target enrichment was achieved through the OneSeq system. Data analysis was conducted using Agilent’s SureCall software. For the cancer-focused and targeted driver mutations analysis, the Ion Torrent AmpliSeq Colon Cancer Panel (AmpliSeq Library 96LV Kit 2.0, ThermoFisher) was employed as an alternative to WES and the OneSeq method. Library preparation, quality control, sequencing, and data analysis were performed as per manufacturer instructions.

### Immunohistochemistry

IHC of patient tissues (see patient-derived organoids methods above) started with immediate fixation of tumor or normal tissue fragments from surgical resections, using 10% neutral buffered formalin. After 24 hours of fixation, specimens were transferred to 70% ethanol, and then processed into paraffin blocks within 1 week. For PDO sections, organoids were grown and treated, then isolated from Matrigel domes using Cell Recovery Solution (Corning #354253).

PDOs were fixed with 4% PFA, embedded in 1% agarose, then processed into FFPE blocks. 5 µm thick sections were subjected to standard IHC. Visualization was with ImmPACT DAB Substrate Kit (Vector Laboratories #SK-4105). Images were acquired using the Albert Einstein Analytical Imaging Facility (AIF) slide scanner (3DHistech Pannoramic 250 digital slide scanner, CMOS detector PCO edge 4.2). See Supplementary Table 2 for primary antibodies and conditions.

### CDK9 expression scoring

Scoring was conducted by a board-certified clinical pathologist specializing in gastrointestinal malignancies (P.Patil). Adjacent H&E slides were used as a reference for each CDK9 stained slide, to identify malignant epithelial cells. Slides were scored and categorized into the following groups: 0 (no nuclear staining), 1+ (<10% positive), 2+ (10%-25% positive), 3+ (26%-50% positive), 4+ (51%-75% positive), and 5+ (>75% positive).

### Statistical analyses

Statistical analyses were carried out using GraphPad Prism, v9.3.1. For CDK9 IHC expression, comparison of matched normal colonic versus CRC tissues was tested using Wilcoxon matched-pairs signed rank test with the following parameters: two-sample matched pairs, two tails. Unmatched testing of the same cohort was performed using the Mann-Whitney test with the following parameters: assumed random selection and independence of samples, assumed non-normal distribution. Survival of TCGA CRC cohorts were analyzed using Kaplan-Meier analysis, with cohorts defined as above or below the CDK9 expression median, and assuming independence of survival times of individuals in each group. For cell culture experiments, p values were calculated using the Student’s t-tests, conducted as unpaired, two-group, two-tails tests, assuming normal distribution and equal variance. Mean ± SD are reported in the figures. Differences were considered statistically significant if p < 0.05. For details on statistical analysis of RNAseq and DNAseq, see Supplementary Methods.

## RESULTS

### CDK9 overexpression correlates with worse survival in CRC

Numerous studies have demonstrated overexpression of CDK9 in solid tumors^20–23^, as well as methods to target CDK9 or adjacent transcriptional activator molecules for cancer therapy^13–15,24–27^. To investigate the expression of CDK9 specifically in CRC, we first analyzed transcriptomic and proteomic data from publicly available databases. At the protein level, CDK9 is overexpressed in CRC when compared to normal colonic tissue (Fig 1A). We next investigated protein expression levels of CDK9 within the tissues from our CRC patients at MECCC (Supp Fig S1A). We found CDK9 protein to be overexpressed in CRC tumor cells when compared to matched normal colonic epithelial cells (Fig 1B, Supp Fig S1B). Interestingly, the expression of CDK9 was heterogeneous between different CRC cells within each patients’ tumor, which potentially reflects intratumor transcriptional heterogeneity. Quantification by histologic scoring demonstrated a significant increase in CDK9 protein in both the overall unpaired comparison of CRC versus matched normal colon (Fig 1C) as well as in the paired comparison (Fig 1D). Furthermore, analysis of CRC cohorts on The Cancer Genome Atlas revealed that overexpression of CDK9 is associated with worse survival in CRC patients of all stages (Fig 1E) including metastatic disease (Fig 1F)^28^. In summary, we found that CDK9 is overexpressed in CRC when compared to matched normal colon tissue and that CDK9 expression levels correlate with worse survival.

**Figure 1.**
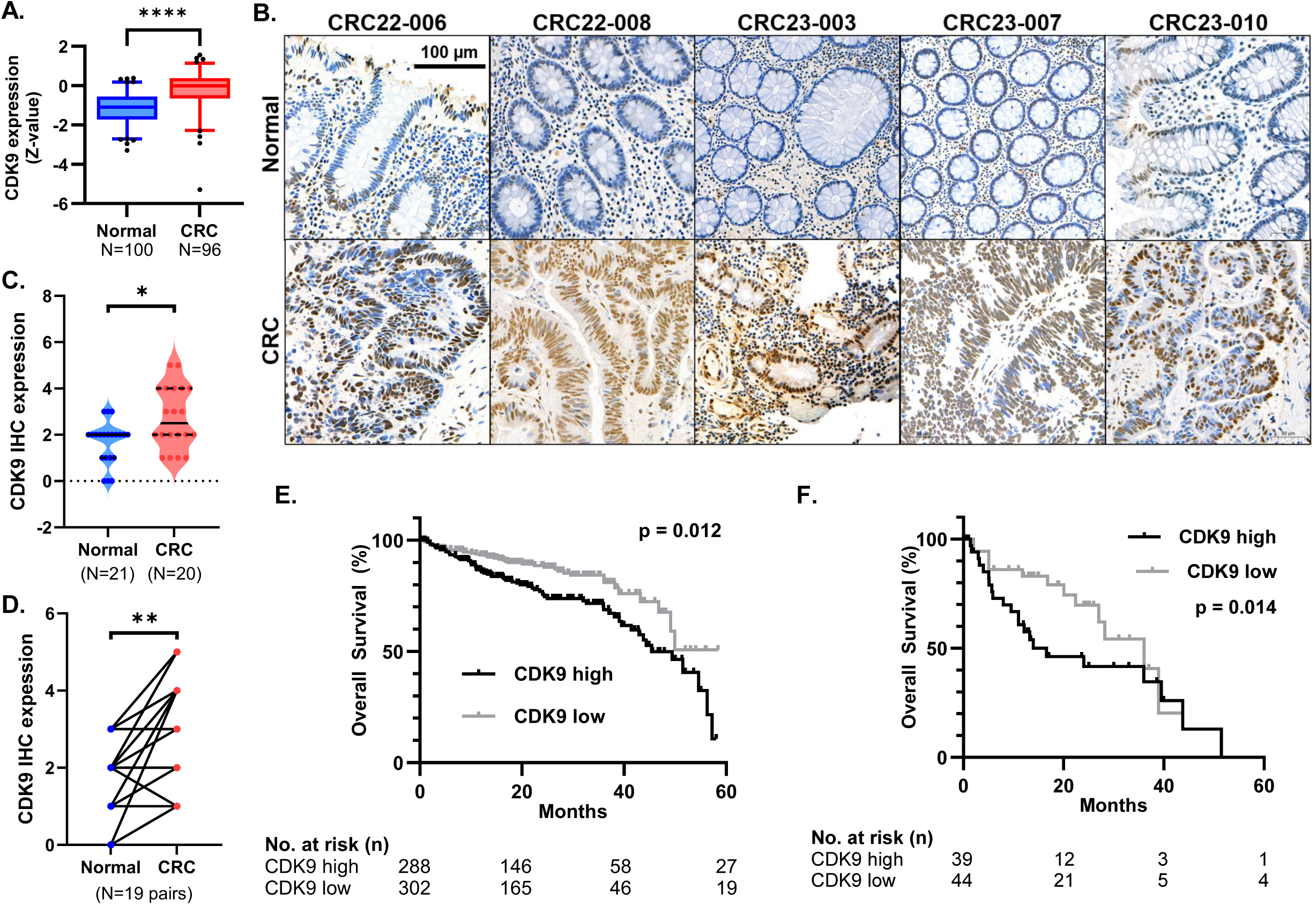
CDK9 is overexpressed in CRC and associated with worse outcomes. A) CDK9 protein expression in CRC versus normal colon tissues from CPTAC proteomic database. ****: p<0.00005. B) Representative images of CDK9 protein expression by IHC in matched CRC and normal colon epithelial tissues from MECCC. C) Quantification of CDK9 protein expression by H-score in matched CRC and normal colon epithelial tissues from MECCC, compared using non-parametric testing. *: p<0.05. D) Comparison of CKD9 protein expression change in matched normal colon and CRC tissue, using paired/parametric testing. *: p<0.05. E) Overall survival of CRC patients of all disease stages as measured by Kaplan-Meier analysis, comparing greater than versus less than median *CDK9* mRNA expression from TCGA. F) Overall survival of stage IV CRC patients as measured by Kaplan-Meier analysis, comparing greater than versus less than median *CDK9* mRNA expression from TCGA.

### CDK9 inhibitors potently suppress CRC cell growth and survival

CDK9 can potentially be targeted by small molecule inhibitors. Rahaman *et al* previously demonstrated that treatment of CRC cells with the CDK inhibitor, CDKI-73/LS-007, can inhibit CRC cell growth and survival, and suppress cancer signaling pathways^25^. Importantly, CDK9 knockdown by siRNA was able to phenocopy small molecule inhibition. Shen *et al* previously demonstrated the *in vitro* efficacy of the CDK2/9 inhibitor dinaciclib in CRC cell lines^20^. However, many CDK inhibitors are nonspecific and can induce off-target effects through inhibition of other kinases^29^. Several newer, CDK9-specific inhibitors have recently been characterized and are being actively developed in early phase clinical trials. We selected three novel CDK9 inhibitors with strong clinical potential to focus our studies on: AZD4573, enitociclib (formerly known as BAY1125152/VIP152), and NVP-2. Each of these CDK9 inhibitors were found to be highly selective for CDK9 versus other kinases, and they have each demonstrated promising anti-tumor activity *in vitro* and *in vivo*^17,24,30^. Ongoing clinical trials to test the safety and efficacy of these drugs are currently focused on hematologic malignancies (NCT04630756, NCT05371054). Thus, the activity and mechanisms of action of these drugs in gastrointestinal cancers, and specifically CRC, are not known.

Using a panel of human CRC cell lines, we found that all three CDK9 inhibitors potently suppressed CRC cell growth and survival in a treatment assay (Fig 2A,B) as well as a colony formation assay (Fig 2C,D) across a wide range of concentrations. We investigated drug activities in such a genomically diverse panel of CRC cells to capture the heterogeneity that exists in CRC. While there were some variations between the potency of each drug and cell line, we generally observed that AZD4573 and NVP-2 were the most potently cytotoxic drugs. Enitociclib was approximately one order of magnitude less potent than the other two drugs, but we still observed CRC cell 50% inhibition concentrations (IC_50_) in the nanomolar range (Fig 2B). While clinical pharmacokinetic (PK) data on AZD4573 and NVP-2 are not yet published, the single dose PK of enitociclib has been studied^30^. The IC_50_ of enitociclib against each of our PDOs was in the nanomolar range, which was achievable with a single dose of enitociclib in humans. Thus, CDK9 inhibitors seem to be highly active in suppressing CRC PDOs at clinically relevant concentrations.

**Figure 2.**
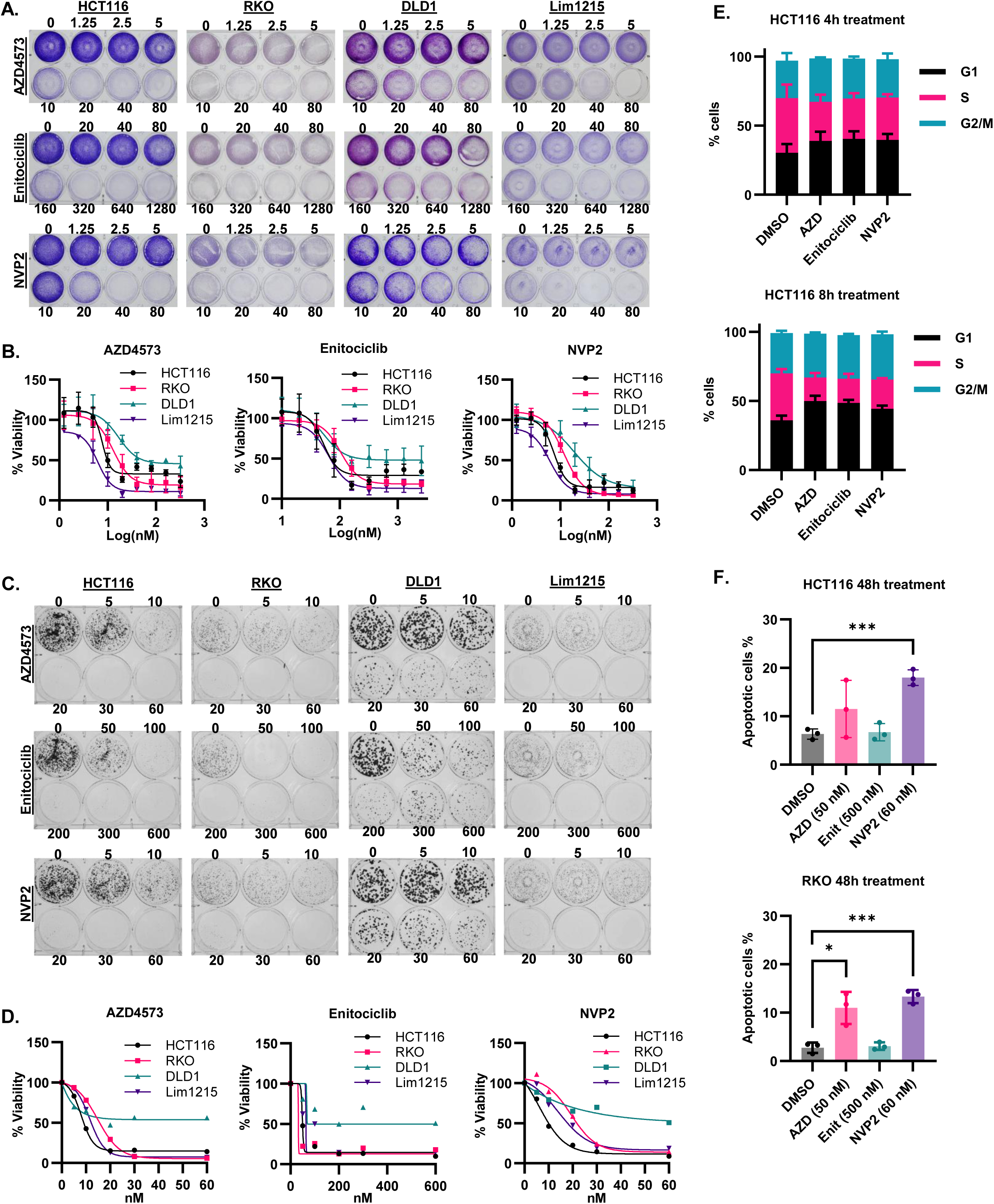
CDK9 inhibition suppresses CRC growth and survival *in vitro*. A) Cell growth assays visualized by crystal violet in cultures of indicated CRC cell lines after 48 h of treatment by CDK9i’s. Drug concentrations (in nM) indicated over each well. B) Cell viability quantification of indicated cell lines treated with CDK9i’s, including logarithmic regression fitted curves. C) Colony formation assays of indicated CRC cell lines after 24 h of treatment by CDK9i’s. Drug concentrations (nM) indicated over each well. D) Colony formation assay quantification of indicated lines treated with CDK9i’s. E) Summary of cell cycle profiles of HCT116 cells treated with each CDK9 inhibitor for 4 or 8 hours (AZD 50 nM, enitociclib 500 nM, NVP2 60 nM). Mean and standard deviation of 3 replicates is shown. F) Change in % apoptotic cells in HCT116 and RKO cells treated with each CDK9 inhibitor at indicated concentrations for 48 hours, as measured by Annexin V and PI staining followed by flow cytometry. Annexin V positive cells were pooled (both late and early apoptosis cells) for this quantification. Mean and standard deviation of 3 replicates shown. *: p<0.05. ***: p<0.0005.

Common causes of decreased cancer cell survival and proliferation include decreased cell cycling, and increased apoptosis. Cell cycle analysis is relevant to this class of inhibitors, because of the potential activity against Cyclin-dependent kinase 4/6 (CDK4/6) inhibitors^31^. Analysis of cell cycle profiles in CRC cells treated for 4 h and 8 h demonstrated only a modest shift in the population of cells, with slightly more cells being in the G1 phase when treated with CDK9 inhibitors. However, this change was not substantial, and a significant proportion of cells remained in the S and G2/M populations (Fig 2E, Supp Fig S2A). We purposefully examined cell cycle profiles at timepoints prior to 24 hours, to determine if cell cycle inhibition is an early or primary mechanism of action by these CDK9 inhibitors, rather than a secondary downstream effect. The limited impact on cell cycle profiles at both 4 h and 8 h seems to suggest that CDK9 inhibitors do not directly affect cell cycle progression. We also assessed the degree of apoptosis in CRC cells treated with CDK9 inhibitors and determined that treatment caused a significantly higher percent of cells to undergo cell death (Fig 2F, Supp Fig S2B). Our results suggest that both apoptosis and cell cycle inhibition play some role in CDK9 inhibitor activity against CRC, but the induction of apoptosis appears to be far more significant.

### CDK9 inhibitors potently suppress the growth and survival of CRC PDOs

To further validate the therapeutic effects of CDK9 inhibitors *in vitro*, we treated a panel of patient-derived organoids (PDOs) established from CRC tumor specimens obtained from patients treated at the MECCC. We established six novel PDOs from six different patients diagnosed with colon or rectal cancer at our Cancer Center, comprising distinct clinical and genomic features (Supp Fig S3). The tumor morphology between pairs of tumor tissue and PDOs were well-matched (Fig 3A). Furthermore, there was significant overlap in the genomic and transcriptomic profiles of the tumors (Fig 3 B,C, Supp Fig S4,S5). These results suggest that our PDO models are biologically representative models of their parental carcinomas.

**Figure 3.**
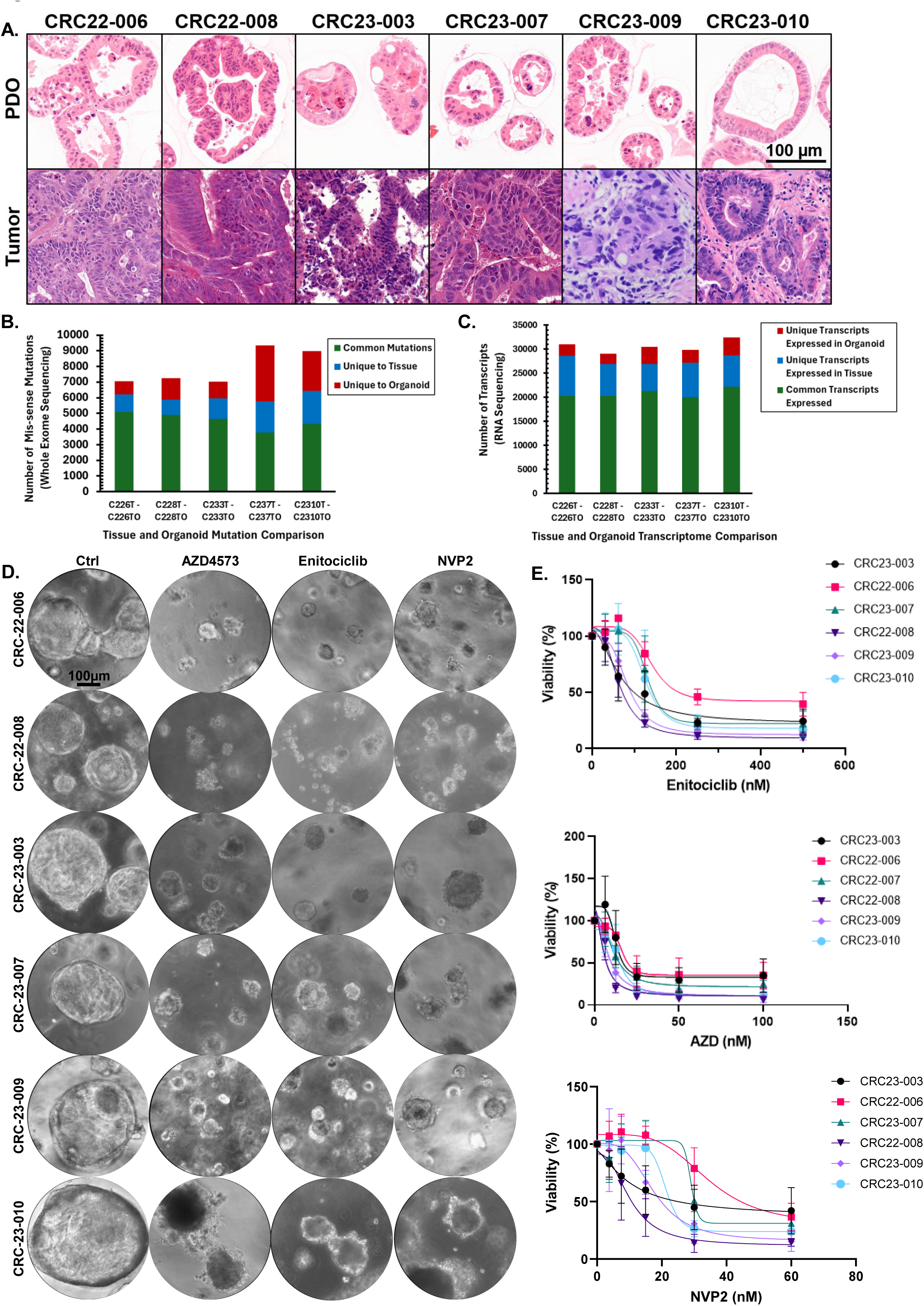
CDK9 inhibition therapeutically suppresses CRC PDO growth. A) Representative images of PDO and parental CRC tissue, stained with H&E. B) Comparison of specific single nucleotide mutations detected in PDOs and matched parental CRC tissues, as measured by whole exome sequencing and referenced to normal wild type hg38. C) Comparison of specific RNA transcripts expressed in PDOs and matched parental CRC tissues, as measured by RNAseq. D) Representative photos of CRC PDOs treated with CDK9i’s, as compared to DMSO control, for 120 hours. Concentrations used: AZD4573 = 50 nM, enitociclib = 500 nM, NVP2 = 60 nM. E) Cell viability quantification of indicated PDOs treated with CDK9 inhibitors, including logarithmic regression fitted curves.

The PDO models used for treatment in this study are heterogeneous in their clinical, molecular, and histopathologic features. Yet nearly every PDO tested was highly sensitive to drug treatment by all three CDK9 inhibitors (Fig 3D), with IC_50_’s in a similar order of magnitude as that found in the CRC cell line treatments (Fig 3E, 2B). One PDO tested, CRC-22-006, was notably less sensitive to treatment than the others, but still had drug sensitivity with IC_50_ in the nanomolar range. Since CDK9 is broadly overexpressed in CRC and is the direct target these inhibitors, we tested for correlation between CDK9 expression levels and sensitivity to CDK9 inhibitors. Although AZD4573 and enitociclib efficacy was not significantly correlated with CDK9 expression, we found that greater NVP2 efficacy was significantly correlated with higher CDK9 protein expression (Supp Fig S6). Overall, our findings suggest that novel, CDK9-specific small molecule inhibitors can potently inhibit the growth and survival of human CRC cell lines and PDOs *in vitro*.

### CDK9 inhibitors suppress multiple downstream cancer signaling pathways including MAPK

We next used both unbiased and targeted approaches to investigate the mechanisms of action of CDK9 inhibitor treatments on our CRC models. For our mechanistic studies, we chose to use inhibitor concentrations within 1- to 2-fold of the IC_50_ for each molecule, to avoid potentially using clinically unrealistic concentration ranges and to preserve the translational relevance of our findings. Since CDK9 is a transcriptional activator, we hypothesized that treatment with CDK9 inhibitors would exert their anti-tumor effect by suppressing the transcription of oncogenes. To study this, we first performed RNAseq on both HCT116 and RKO cells treated by CDK9 inhibitors at multiple concentrations. As expected, we observed an overall change of the CRC cells’ transcriptome due to treatment with each drug, with the higher dose of each drug causing more transcriptional changes (Supp Fig S7A). Interestingly, rather than seeing global downregulation of most transcripts due to CDK9 inhibitor treatment, we observed that more genes were upregulated with high dose CDK9 inhibitor treatment. A broad view of the transcriptional changes by principal component analysis demonstrated that treatment with different drugs led to similar transcriptional profiles, and these transcriptomes were drastically different from those of control or untreated CRC cells (Supp Fig S7B). Overall, our RNAseq results suggest that treatment with CDK9 inhibitors significantly alters the CRC cell transcriptome in a similar fashion as one another, concordant with their shared mechanism of suppressing CDK9.

To further elaborate on the downstream targets affected by CDK9 inhibitors, we conducted Parametric Analysis of Gene Set Enrichment (PAGE) on the RNAseq data (Supp Fig S8)^32^. Several key biological and oncologic pathways were identified as being significantly suppressed by CDK9 inhibitor treatment. Notably, the gene transcription pathway was found to be one of the most significantly suppressed pathways across all treatment conditions, consistent with the primary mechanism of CDK9 inhibitors in suppressing transcription. Our pathway analysis also uncovered significant suppression of gene sets corresponding to colorectal cancer, MAPK signaling pathway, and mTOR signaling pathway at the RNA level. Due to the importance of MAPK signaling pathway in driving CRC carcinogenesis^33^, we chose to focus on this pathway for further investigation. Detailed assessment of MAPK pathway genes suggested that transcripts of several canonical genes were indeed suppressed by CDK9 inhibitor treatment (Fig 4A,B). We further validated this finding using RT-qPCR and found several MAPK pathway genes suppressed at the mRNA expression level by CDK9 inhibitor treatment (Fig 4C, Supp Fig S9,10). Interestingly, our PCR results suggested that at the mRNA level, *EGFR*, *KRAS*, and *BRAF* were more consistently suppressed as compared to more downstream pathway genes such as *MEK* and *ERK* (Supp Fig S10).

**Figure 4.**
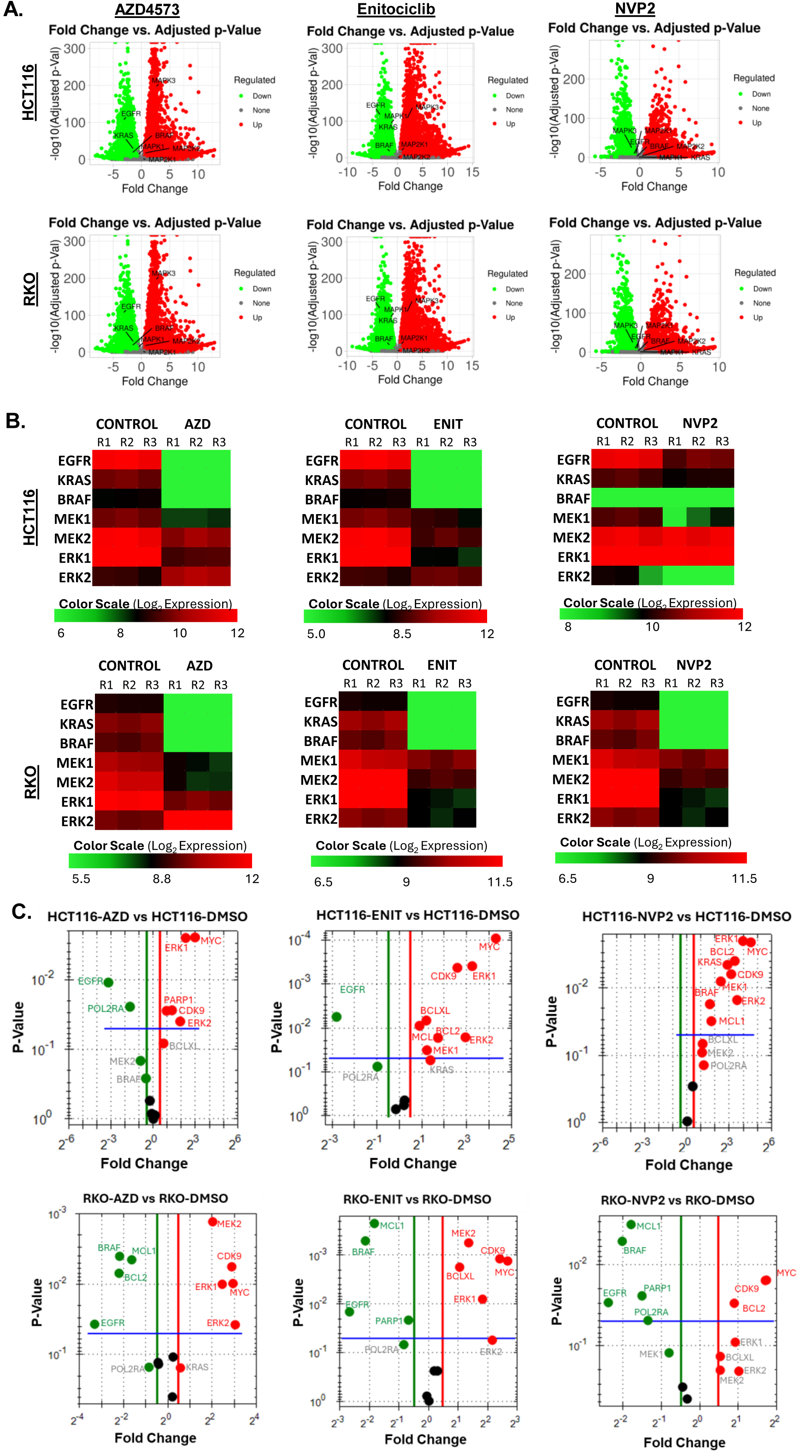
Transcriptomic analysis of CRC cells treated with CDK9 inhibitors. A) Volcano plots of total transcriptome change, based on RNAseq, in CRC cells treated with CDK9 inhibitors for 24 h. Individual transcripts of several MAPK signaling targets are represented. B) Heatmap of transcriptional changes seen in RNAseq of MAPK genes of CRC cells treated with CDK9 inhibitors for 24 h. C) RT-qPCR expression levels of MAPK genes in CRC cells treated with CDK9 inhibitors for 24 h, as compared to DMSO control treatment. Concentrations used: AZD4573 = 50 nM, enitociclib = 500 nM, NVP2 = 60 nM.

While not the focus of this study, we also assessed the mRNA levels of the canonical CKD9 targets, such as *MYC* and *MCL1*. These genes are believed to be particularly dependent on CDK9 activity because they generate rapidly degraded transcripts, and therefore require active transcription to maintain high levels of mRNA transcript^23^. Our PCR data suggest that *MYC* mRNA was not significantly suppressed by any of the drugs we tested (Supp Fig S10). In contrast, *MCL1* mRNA was significantly suppressed in several of our treatment experiments (Supp Fig S10). These results speak to potential differences between how CDK9 and CDK9 inhibitors might be functioning in CRC as compared to other tumors^14,34^.

To further investigate the transcriptional modulation caused by CDK9 inhibitor treatment, we performed RT-qPCR on our PDOs after treatment. Consistent with our CRC cell line results, we found numerous CDK9 downstream targets to be transcriptionally suppressed by CDK9 inhibitor treatment (Supp Fig S11). Overall, our analysis of the transcriptional changes caused by CDK9 inhibitor treatment suggested a broad inhibition of MAPK signaling pathway genes, along with suppression of *MCL1*.

Our transcriptional analysis strongly suggests that CDK9 inhibitors caused changes in the expression of MAPK signaling genes. However, the MAPK pathway depends on the MAPK proteins to signal and drive cancer proliferation and survival. Therefore, we next assessed the expression of MAPK signaling proteins, as well as other protein targets that could be affected by CDK9 inhibitor treatment. Immunoblotting of key MAPK pathway proteins demonstrated significant reduction in expression of multiple targets along the signaling pathway, across multiple cell lines and across the different drugs tested (Fig 5A,B, Supp Fig S12). Of note, assessment of CDK9 expression and RNA Pol II expression after CDK9 inhibitor treatment demonstrated a decrease in overall RNA Pol II expression, as well as phosphor-Ser2 of RNA Pol II (Supp Fig S13). Quantification of the cell death signal, cleaved PARP, demonstrated significant increase in cell death as a downstream mechanism of CDK9 inhibitor treatment (Supp Fig S14A,B). We also observed inconsistent and, in some cases, absent reduction of c-MYC and MCL-1 expression across the CRC cell lines tested (Supp Fig S14A,C). This finding appears to be a unique observation in contrast to the RNA Pol II mechanisms seen in hematologic malignancy studies, where there is more consistent suppression of c-MYC and MCL-1 at both the transcript and protein levels^17^.

**Figure 5.**
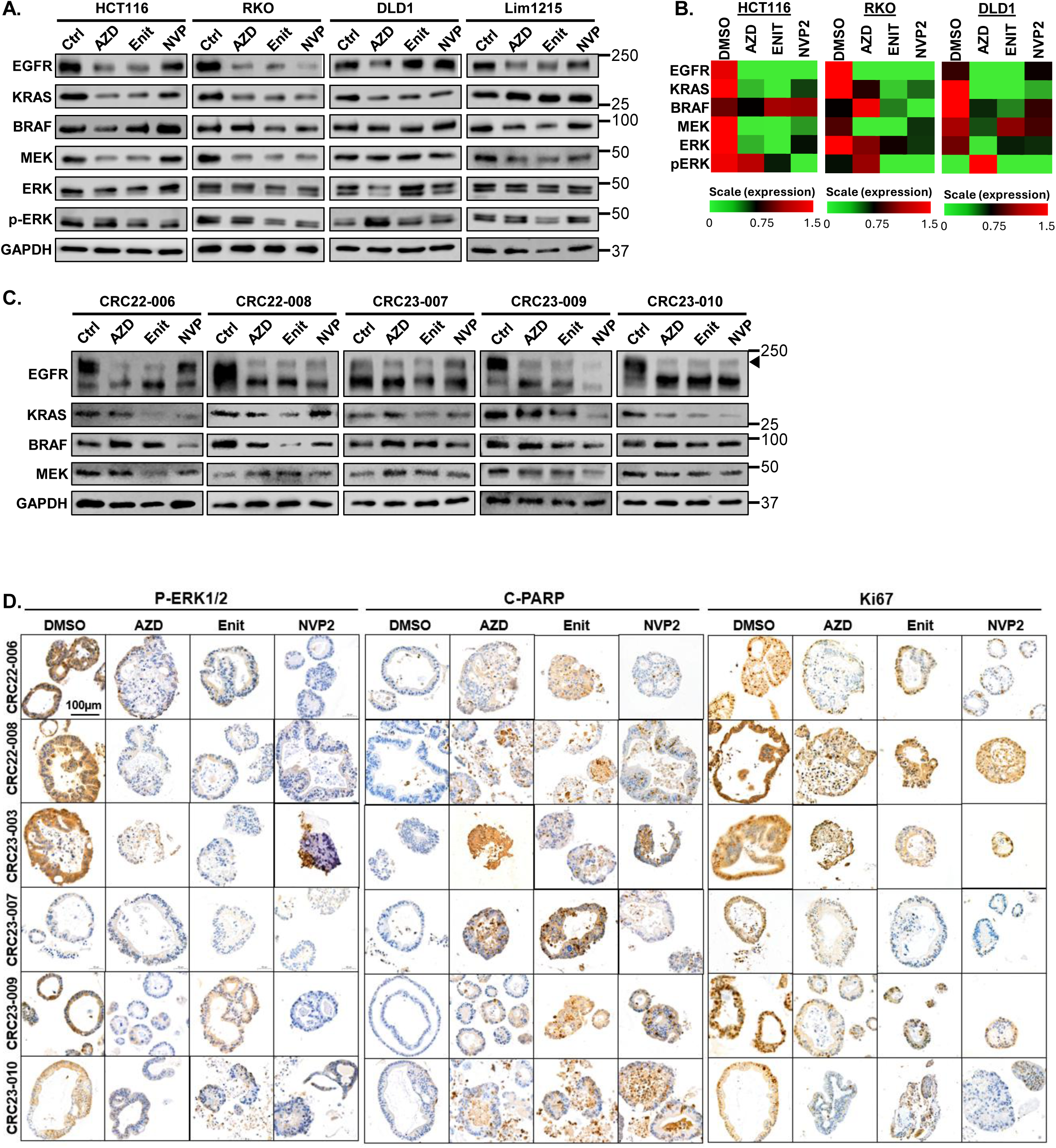
Protein analysis of CRC cells and PDOs treated with CDK9 inhibitors. A) Representative immunoblot of MAPK signaling pathway protein expression in CRC cells treated with CDK9 inhibitors for 24 h. Drug concentrations: AZD4573 = 50 nM, enitociclib = 500 nM, NVP2 = 60 nM. B) Heatmap representation of immunoblot data shown in 5A. Three replicates were pseudo-quantified by pixel densitometry and averaged. C) Representative immunoblot of MAPK targets in PDOs treated with CDK9 inhibitors for 24 h. D) Representative images of CRC PDOs treated with DMSO or CDK9 inhibitors for 5 days and visualized by IHC with antibodies against indicated proteins. Concentrations used: AZD4573 = 50 nM, enitociclib = 500 nM, NVP2 = 60 nM.

To evaluate and confirm these mechanisms using our patient-derived models, we turned again to our PDOs. Similar to our findings at the transcriptional level, we found significant suppression of several MAPK proteins (Fig 5C,D). As with the CRC cell lines, CRC PDOs also seem to demonstrate decreased RNA Pol II expression in response to CDK9 inhibitor treatment (Supp Fig S15A). In contrast to the cell lines, however, the PDOs did seem to have more significant suppression of c-MYC and MCL-1 because of CDK9 inhibition (Supp Fig S15B).

To address the dynamics of CDK9 inhibitor modulation on MAPK pathway signaling, we studied both the concentration and time dependence of MAPK downstream kinase signaling. At concentrations close to the IC_50_ of each drug, we found significant suppression of p-MEK, p-ERK, and p-RSK (Supp Fig S16A), further suggesting that MAPK signaling suppression is indeed part of the mechanism of CDK9 inhibitors in CRC. We further determined that suppression of MAPK downstream kinase signaling occurs within hours of treatment, indicating that is potentially a primary mechanism of CDK9 inhibition in CRC (Supp Fig S16B). To confirm if the suppression of MAPK kinase signaling is leading to loss of MAP transcriptional output, we analyzed the gene expression changes of MAP network genes from our RNAseq experiment. After CDK9 inhibitor treatment, we observed more MAPK-regulated genes being transcriptionally suppressed than activated (Supp Fig S17). Overall, our findings suggest that CDK9 inhibitors work by shifting the transcriptional landscape of CRC, with significant impacts on multiple cancer signaling pathways and in particular on the MAPK signaling pathway.

### Rational combinations of CDK9 inhibitors and MAPK pathway inhibitors synergize to treat CRC

While CDK9 inhibition has been shown to impact different cancer pathways in a context-dependent fashion, we suspected that the “vertical suppression” of the MAPK signaling pathway may be a unique feature of CDK9 inhibitors in CRC. We hypothesized that suppression of this signaling pathway by CDK9 inhibition could be leveraged therapeutically by combining with targeted agents against MAPK proteins. In our study, we observed that EGFR and KRAS protein expression were already deeply suppressed by CDK9 inhibitor treatment. We reasoned that a less suppressed signaling protein that was also in the MAPK pathway might be a more suitable target for combination treatments. Therefore, we chose to test the combination of MEK inhibitors and CDK9 inhibitors. For MEK inhibitors, we selected the clinically promising drugs cobimetinib and mirdametinib. We found that combined treatment of CDK9 inhibitors and MEK inhibitors was only modestly synergistic against a PDO derived from a patient with MSI-H CRC (PDO CRC-22-008, Fig 6A). However, these same combination treatments were much more potently synergistic against an MSS CRC with a TP53 mutation (PDO CRC-23-007, Fig 6B). In particular, the combination of enitociclib and MEK inhibitor was strongly synergistic across a wide range of drug concentrations. Overall, our results suggest that CDK9 inhibitors and MEK inhibitors can synergistically suppress CRC tumor growth and survival. These findings support the hypothesis that CDK9 inhibitors could be combined with direct inhibitors of MAPK signaling proteins in order to leverage the unique ability of CDK9 inhibitors to suppress MAPK protein transcription.

**Figure 6.**
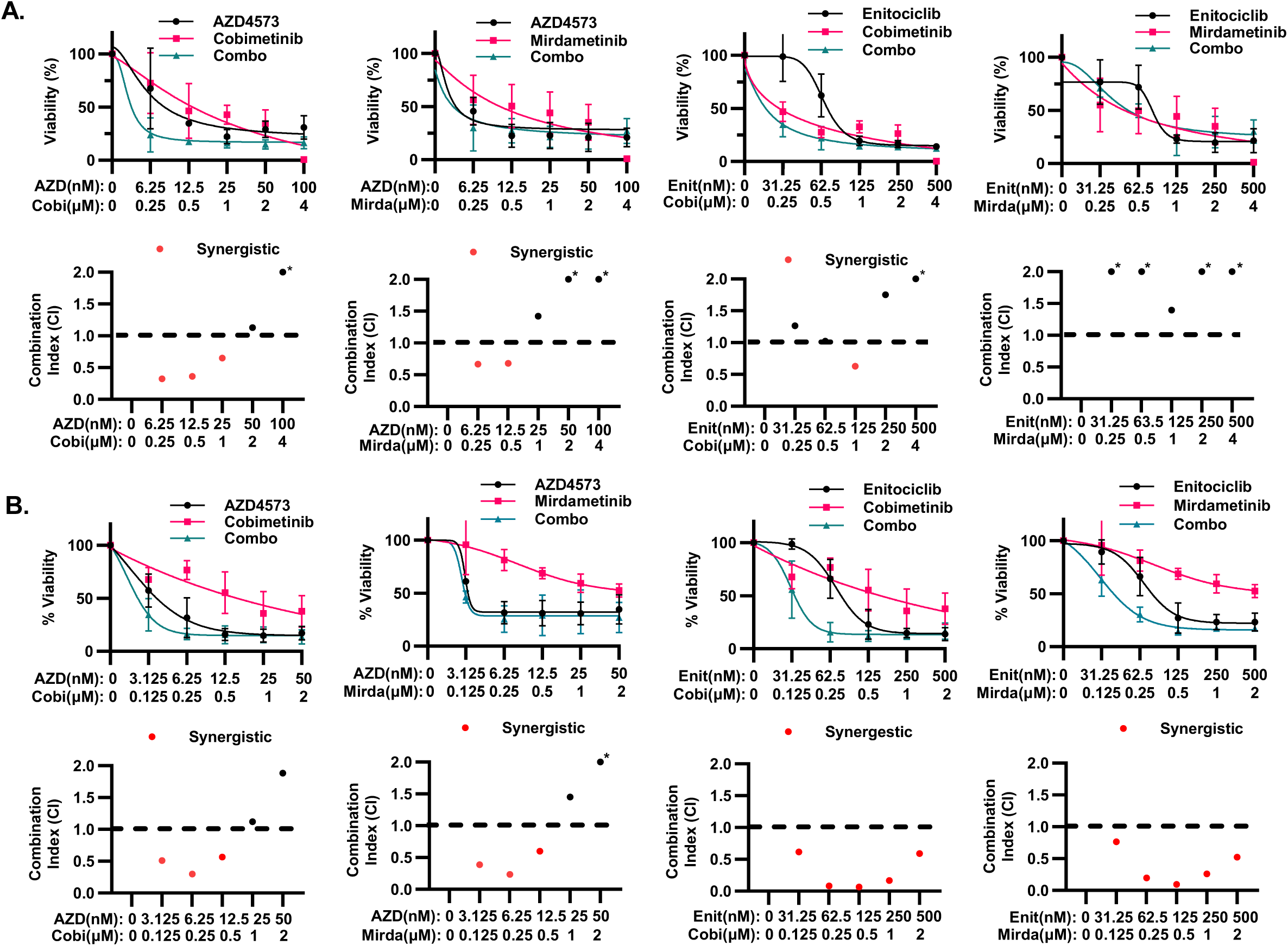
Treatment of CRC PDOs with the combination of MEK inhibitors + CDK9 inhibitors. A) Cell viability of CRC-22-008 treated with either mirdametinib or cobimetinib combined with CDK9 inhibitors for 5 days. Synergy scores are shown directly below the viability graphs. B) Cell viability of CRC-22-007 treated with either mirdametinib or cobimetinib combined with CDK9 inhibitors for 5 days. Synergy scores, as calculated by the Chou-Talalay model, are shown directly below the viability graphs.

## DISCUSSION

Recent development of more specific CDK9 inhibitors have re-energized this field within cancer therapeutics^17,24,30,35,36^. Yet, there are still few studies detailing the activity and mechanisms of action by these molecules in solid tumors, especially CRC. To our knowledge, our study is the first to provide such a comprehensive analysis of AZD4573, Enitociclib, and NVP2 in CRC. We found that CDK9 inhibitors are highly potent inhibitors of CRC growth and survival. An important finding in our study is that low nanomolar concentrations of CDK9 inhibitors can suppress CRC growth and survival. For AZD4573 and NVP-2, we are not able to draw direct comparisons to human drug exposure, due to the limited clinical data available. However, we were able to compare our experimental concentrations of enitociclib with published clinical PK data^30^. We found the IC_50_ of enitociclib to be consistently below 200 nM, well within the range of drug exposure seen in the phase 1 study of enitociclib. Therefore, the concentrations of CDK9 inhibitors used in our study are likely within a clinically realistic range. Our mechanistic data revealed both expected alterations, such as suppression of RNA Pol II and transcription, as well as unexpected effects, such as the suppression of MAPK and PI3K/AKT pathways.

Subsequently, we were able to capitalize on the finding of MAPK suppression by demonstrating synergy between CDK9 inhibitors and MEK inhibitors in CRC. Thus, our study highlights the therapeutic potential of CDK9 inhibitors not only as monotherapy, but also as rational combination therapies. The data generated herein can be used to generate many new hypotheses about novel combinations incorporating CDK9 inhibitors with realistic clinical potential.

Another highlight of our study is the use of CRC tissue specimens and models generated from our unique patient population. The MECCC primarily serves the Bronx, NY catchment area, which is heavily enriched with underrepresented groups^37^. We first conducted an independent analysis of CDK9 expression in CRC and normal colonic tissues derived from our unique patient population. Our cohort’s racial and ethnic diversity is highly consistent with our hospital catchment area’s overall patient diversity. Our finding that CDK9 is highly expressed in CRC as compared to colonic mucosa is consistent with previous findings in different cohorts and suggests that CDK9 acts as a CRC driver across diverse patient groups. We also established several new PDOs from CRC tissues specimens in our biobank. Our study included rigorous genomic, transcriptomic, and histologic comparison of the parental tissues and PDOs. As documented in numerous prior studies, our PDOs are highly representative of the parental CRC tissues and serve as powerful patient-derived models for the study of cancer biology and therapeutics. The availability of these models to the scientific community will help enrich the diversity of publicly accessible PDOs. Our results can be used to support the rationale for therapeutic testing of these drugs in broadly inclusive groups, which may help rebalance the significant discrepancies in outcomes of patients in underrepresented minorities^38^.

The finding of “vertical suppression” of the MAPK pathway by CDK9 inhibitors was an important and surprising result of this project. Vertical suppression is the inhibition of multiple targets within a single signaling pathway. This method of suppression has been found to overcome adaptive resistance which can often occur when targeting a single molecule within signaling pathways^39,40^. In the case of MAPK driven tumors, many resistance mechanisms appear to converge on re-activation of the EGFR-RAS-RAF-MEK axis^41–46^. Therefore, an orthogonal method of suppressing multiple targets in this signaling pathway, such as through transcriptional suppression, would be clearly beneficial as either monotherapy or combination therapy. Several nuances to our findings are worth further discussion. One possibility, due to the frequent off-target effects seen with small molecule inhibitors, is that MAPK pathway targets such as MEK or ERK could be directly inhibited by the CDK9 inhibitors we studied. We find this to be unlikely, due to the rigorous kinase assays conducted by other groups on each of these molecules. Such assays found either weak or no inhibition of MAPK family signaling molecules by AZD4573, VIP152, and NVP-2^17,24,47^.

Our study begs the question of why CDK9 inhibition would so strongly intersect with MAPK-pathway signaling. While we did not find a clear MAPK-associated molecular alteration in our cell lines and PDO models to explain potential differences in sensitivity to CDK9 suppression, it is possible that an overall dependence of MAPK pathway activation could be a marker of sensitivity to CDK9 inhibitors. For example, the PDO model CRC23-007 was relatively less sensitive to CDK9 inhibitors, particularly NVP2 (Fig 3E). Concurrently, we found this model to show only modest changes in MAPK pathway expression level changes at both the RNA (Supp Fig S11) and protein levels (Fig 5C,D). Additional work is needed to further test the hypothesis that CDK9 inhibitor sensitivity is dependent on cumulative MAPK signaling dependence. It is also important to recognize that even though MAPK signaling modulation was modest in CRC23-007, there was still significant anti-tumor toxicity with CDK9 inhibitor treatment. This could be because many other pathways in CRC are dependent on CDK9-mediated transcriptional upregulation (Supp Fig S8). In contrast to the CDK9 monotherapy results, the combination experiments with MEK inhibitors revealed that CRC23-007 is susceptible to synergistic activity by CDK9 and MEK inhibitors (Fig 6B). This result would suggest that despite the appearance of less dependence of CRC23-007 on MAPK signaling, there may be enough reliance on this pathway to still facilitate this synergistic treatment activity. Overall, these results open the possibility to explore CDK9 inhibitors as upstream modulators of many other oncogenic pathways, as well as novel mechanisms of synergy.

CDK9 inhibitors have been studied most extensively in hematologic malignancies due to the hypothesis that c-MYC and MCL-1, which are CDK9-dependent genes, are also highly upregulated in blood cancers. While we found c-MYC and MCL-1 to be suppressed, in some cases quite strongly, by CDK9 inhibitors, we did not find these to be among the strongest mechanisms induced by CDK9 inhibitor treatment. When taking a pathways approach, we found other cancer signaling pathways to be heavily impacted, such as the MAPK and PI3K/AKT signaling pathways. While the exact reason behind the differences in mechanism are beyond the scope of this study, we hypothesize that CDK9 inhibitors act in a context dependent fashion based on the cancers they are treating. Hematologic malignancies may rely more heavily on c-MYC signaling and MCL-1-dependent apoptosis suppression. CRC carcinogenesis is dominated by MAPK and PI3K signaling^48^. Whichever pathways are most active in a particular type of cancer will be the most “CDK9-addicted” because they will consume as much transcriptional resources as possible to driver oncogenesis. Thus, CDK9 inhibitors could potentially be active against many cancers by acting as suppressors of tumors with high transcriptional activity and dependence. Further study of the biochemical mechanism of transcriptional machinery by CDK9 inhibitors in CRC is ongoing.

Our study has several limitations. We have not yet characterized a mechanism of resistance to CDK9 inhibitors. Our work thus far has revealed that most CRC cells and PDOs are similarly sensitive to CKD9 inhibitor treatment. The most resistant CRC cell line, DLD-1, shares many key cancer driver mutations with other models we report on here, making nomination of hypothetical resistance mutations challenging. Since CDK9 inhibitors are transcriptional inhibitors, potential mechanisms of resistance may lie in the transcriptional or epigenomic profiles of CRC. Work is planned to elucidate these potential mechanisms. Acquired resistance mechanisms can also develop in response to CDK9 inhibitor treatment and warrants further investigation^34^. Another planned future direction is *in vivo* validation of our therapeutic results. For example, *in vivo* validation of CDK9 inhibitors plus MEK inhibitors in patient-derived xenograft models could help to further strengthen the rationale for testing the clinical efficacy of these treatments in CRC patients.

In conclusion, we demonstrate that several recently developed small molecule CDK9 inhibitors are highly efficacious against preclinical models of CRC. We found these treatments to be effective against CRC cells as well as CRC PDOs derived from an urban, minority-rich patient population. Our mechanistic work revealed novel insights about CDK9 inhibitors in CRC, most notably that the MAPK signaling pathway is strongly suppressed. This finding supports a multitude of potential hypotheses about novel combination treatment using CDK9 inhibitors in CRC. Accordingly, we found that CDK9 inhibitors and clinical MEK inhibitors synergize to treat a CRC PDO. Future work may leverage our study’s findings to generate additional supporting data for the rationale of novel clinical trials of CDK9 inhibitors in CRC patients.

## Abbreviations

BRAF: V-Raf Murine Sarcoma Viral Oncogene Homolog B
CDK: Cyclin Dependent Kinase
CDK4/6: Cyclin Dependent Kinase 4/6
CDK9: Cyclin Dependent Kinase 9
CMS: Consensus Molecular Subtypes
c-MYC: Cellular myelocytomatosis oncogene
CRC: Colorectal Cancer
EGFR: Epidermal Growth Factor Receptor
ERK: Extracellular Signal-Regulated Kinases
IC_50_: 50% Inhibition Concentration
KRAS: Kirsten Rat Sarcoma Viral Oncogene Homolog
MAPK: Mitogen Active Protein Kinase
MECCC: Montefiore Einstein Comprehensive Cancer Center
MCL-1: Mixed lineage leukemia 1
mCRC: metastatic Colorectal Cancer
MEK: MAPK Kinase
MSI-H: Microsatellite Instability High
MSS: Microsatellite Stable
mTOR: Mammalian Target of Rapamycin
PAGE: Parametric Analysis of Gene Set Enrichment
PARP: Poly (ADP-ribose) Polymerase
PDOs: Patient Derived Organoids
PK: pharmacokinetic
RNA Pol II: RNA polymerase II
VEGF: Vascular Endothelial Growth Factor

## Acknowledgements

FFPE specimen processing, slide cutting, and H&E staining was conducted by Laura Ramkissoon of the Albert Einstein College of Medicine Histology and Comparative Pathology Facility, which is partially funded by NCI grant P30CA013330. Slide scanning was conducted by Hillary Guzik, Rotem Alon and Vera DesMarais, of the Albert Einstein AIF. The 3DHistech Pannoramic 250 digital slide scanner is housed in the AIF and was purchased with NIH grant SIG S10OD026852-01A1. The AIF is partially funded by NCI grant P30CA013330. Clinical research support was provided by Benjamin Duva, Gabrielle Isom, Chika Ekweghariri, and Rikin Gandhi of the Montefiore Einstein CCTO. The CCTO is supported in part by NCI grant P30CA013330. We acknowledge Jackie Chuen and Kamini Singh for supporting the flow cytometry experiments and Alicia Tiedeman and Stephanie Moldavsky for their support of the patient specimen procurements. The results published here are in part based upon data generated by the TCGA Research Network: https://www.cancer.gov/tcga.

## Author Contributions

M.M.: Experimental planning, data collection, manuscript drafting, manuscript revision.

M.A.B.: Experimental planning, data collection, manuscript drafting, manuscript revision.

T.L.: Experimental planning, data collection, manuscript drafting, manuscript revision.

N.W.: Experimental planning, data collection, manuscript revision.

P.P.: Experimental planning, data collection, manuscript revision.

C.M.: Data collection, manuscript revision.

N.T.: Data collection, manuscript revision.

Z.H.: Data collection, manuscript revision.

J.D.: Project coordination, experimental planning, data collection.

D.Y.G.: Project coordination, experimental planning.

O.W.: Project coordination, manuscript revision.

R.H.: Project coordination, manuscript revision.

K.O.: Project coordination, manuscript revision.

M.Q.: Project coordination, manuscript revision.

E.C.: Project oversight, manuscript revision, funding.

C.K.: Project oversight, project coordination, experimental planning, data collection, manuscript drafting, manuscript revision, funding.

## Ethics Approval and Consent to Participate

All patient specimens were collected from patients enrolled in the MECCC Biobank (IRB# 2021-13730). The study protocol was approved by the Einstein College of Medicine Institutional Review Board. All patients provided written informed consent to participate in the Biobank. All specimens were obtained at minimal risk, after patients had undergone tissue removal using standard of care clinical procedures.

## Data Availability

All data in this manuscript are available upon reasonable request. Genomic and transcriptomic data will be submitted to appropriate databases once the manuscript is in press. Patient-derived organoids will be submitted to appropriate biobanks once the manuscript is in press.

---

**Supplementary Figure S1.**
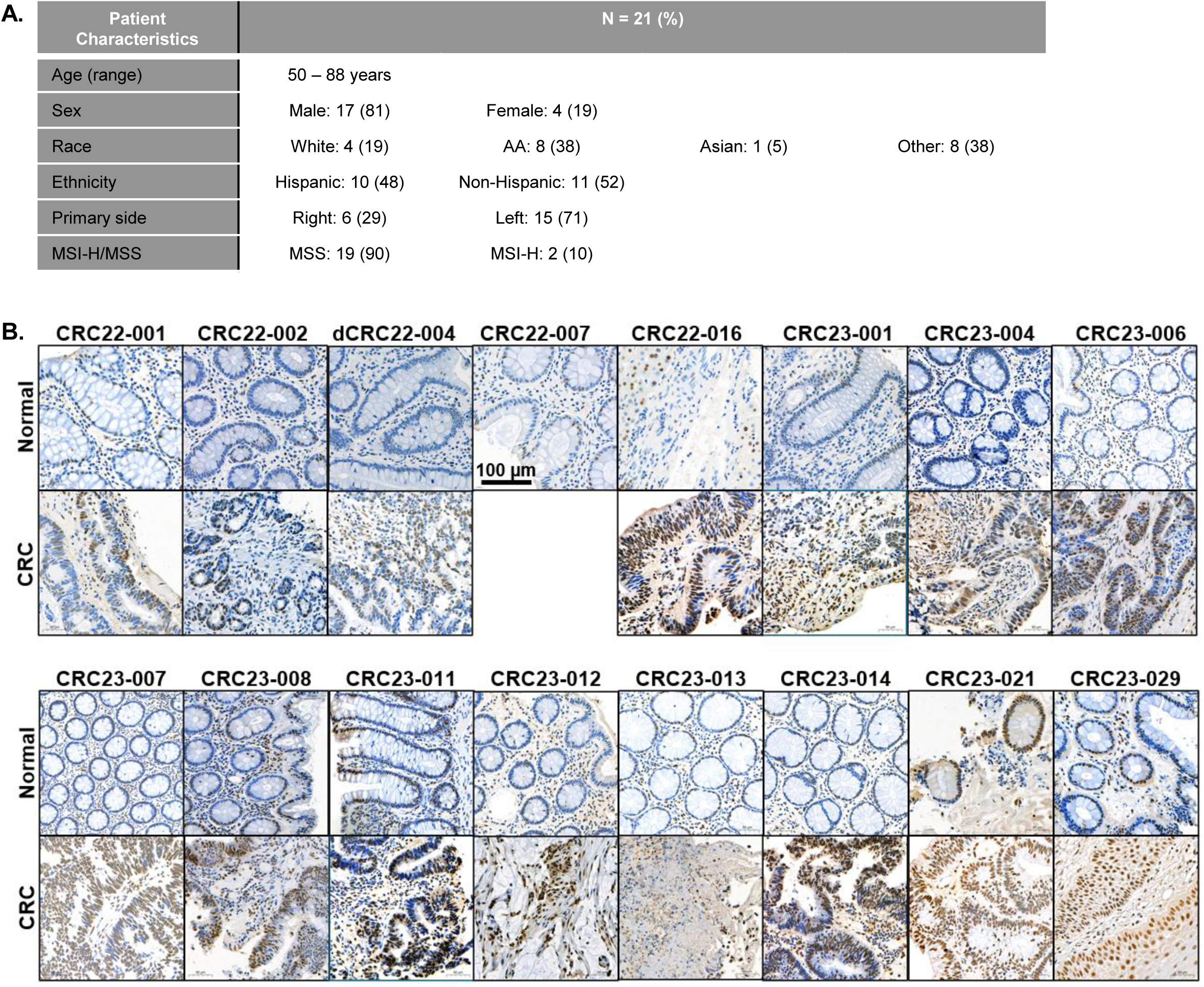
A) Clinical characteristics of MECCC patients included for CDK9 expression analysis. B) Additional representative images of CKD9 IHC stained tissue sections from MECCC.

**Supplementary Figure S2.**
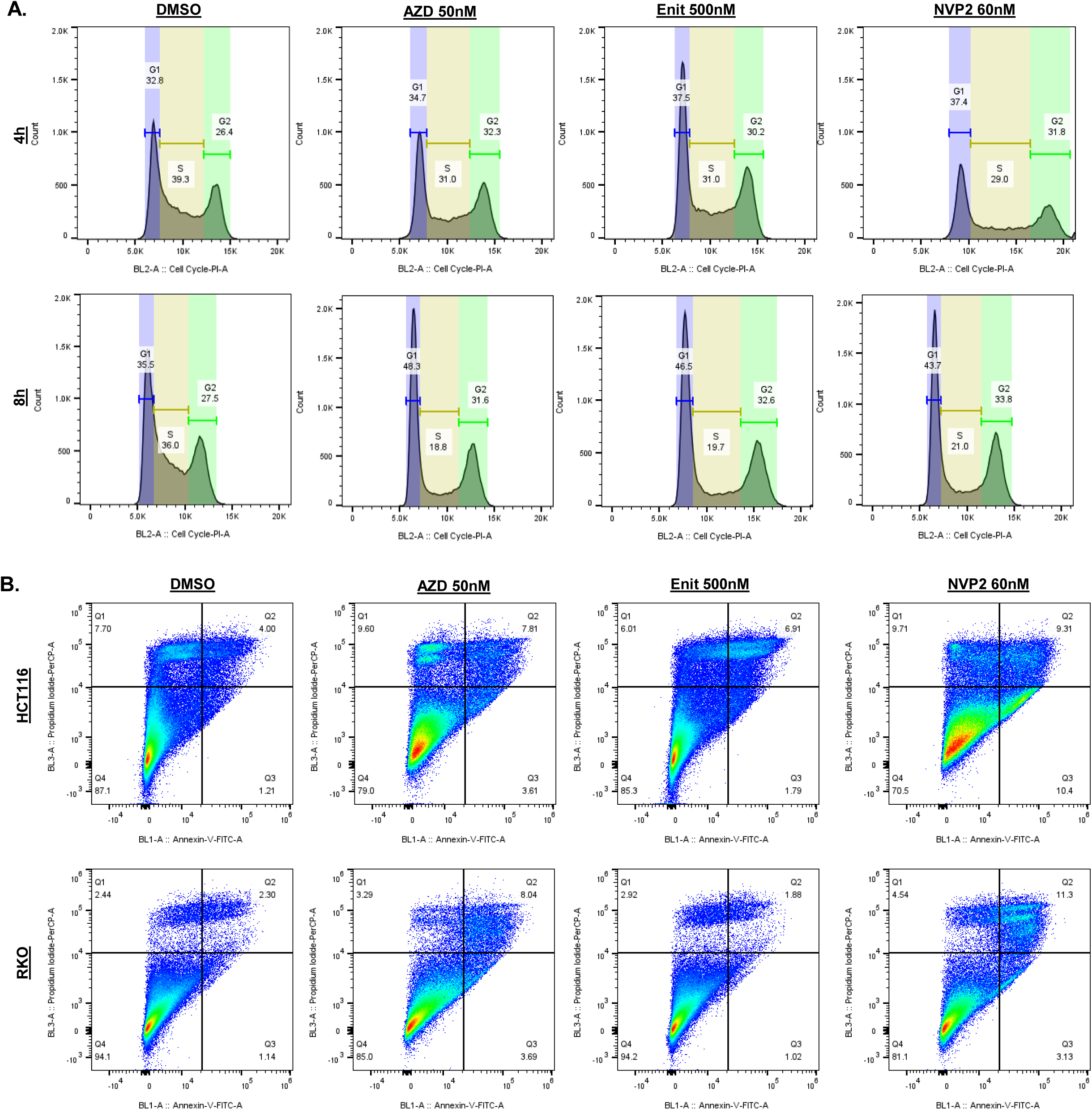
A) Representative cell cycle profiles with fitted curves for HCT116 cells treated with CDK9i’s for 4 h and 8 h., respectively. B) Representative flow cytometry plots of CRC cells treated with CDK9i’s for 48 h and stained with Annexin V and PI.

**Supplementary Figure S3.**
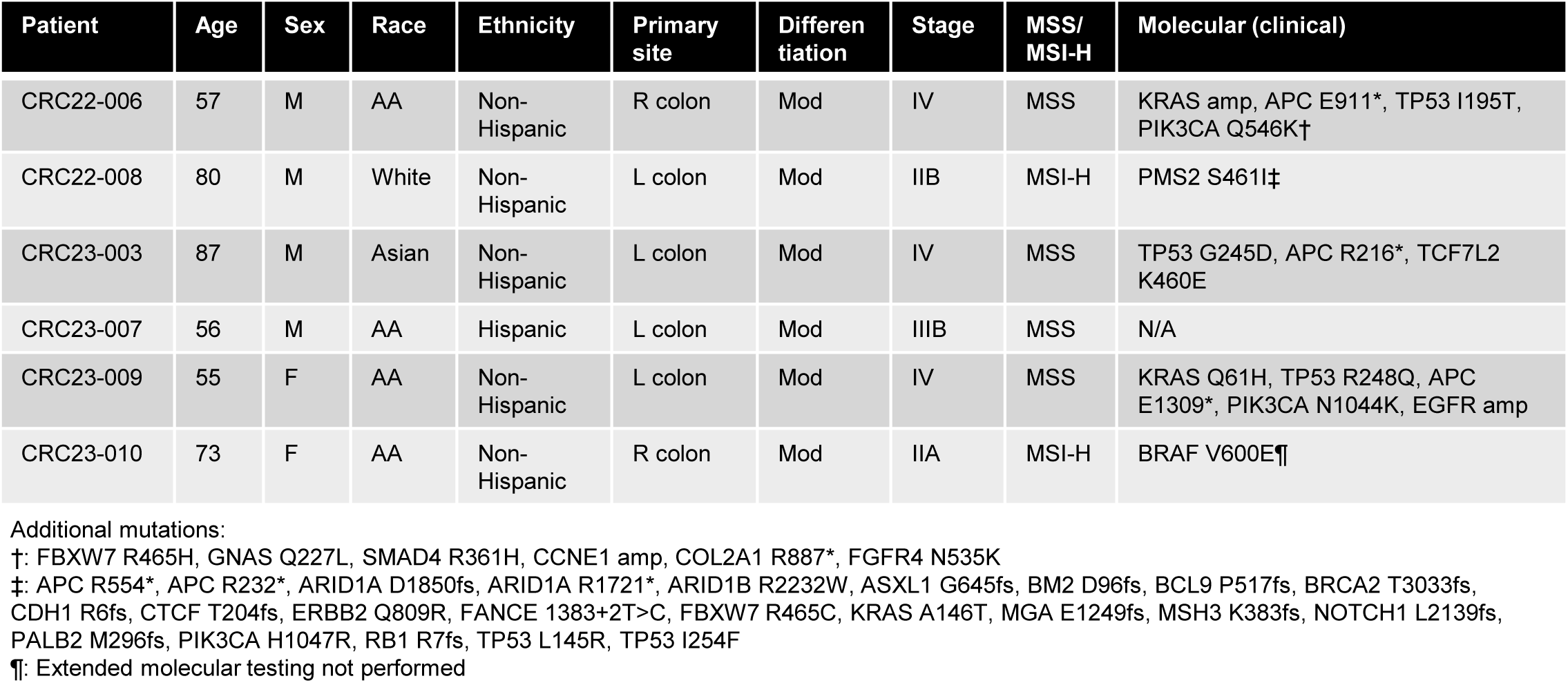
Clinicopathologic features of each PDO used in this study.

**Supplementary Figure S4.**
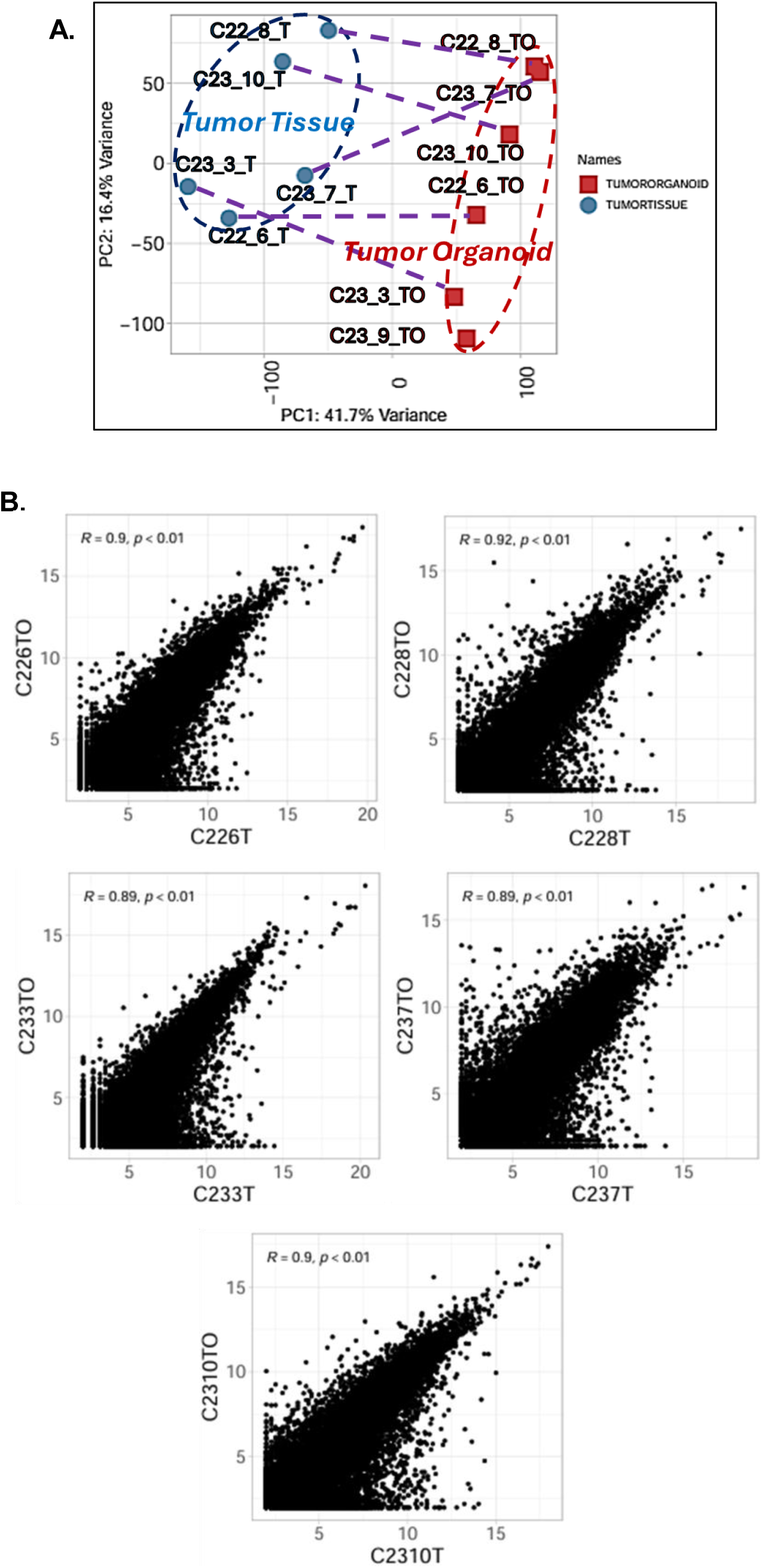
A) 2-dimensional principal component analysis of the transcriptome of the PDOs and their matched parental CRC tissues. Dotted lines connect the matched pairs of tissues/models. B) Spearman correlation between the expression levels of individual gene transcripts in the PDOs versus matched parental CRC tissues, normalized to housekeeping gene.

**Supplementary Figure S5.**
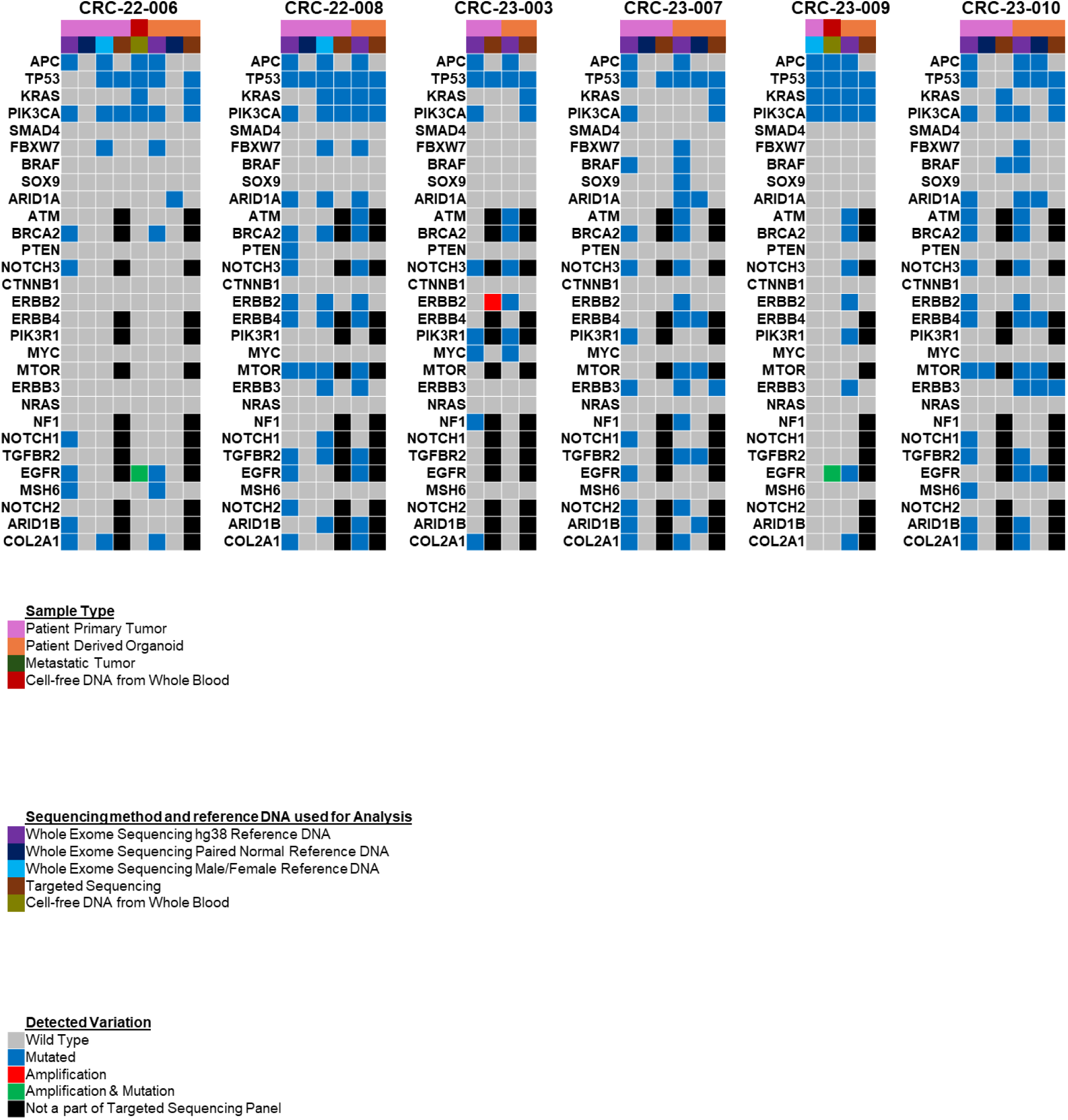
Cross-referenced analysis of gene mutations in cancer driver genes, as assessed by WES or targeted panel sequencing. WES analysis was performed in two ways, one was by referencing a standard hg38 genome, and the second was by referencing the paired normal colon tissue WES sequence as the reference.

**Supplementary Figure S6.**
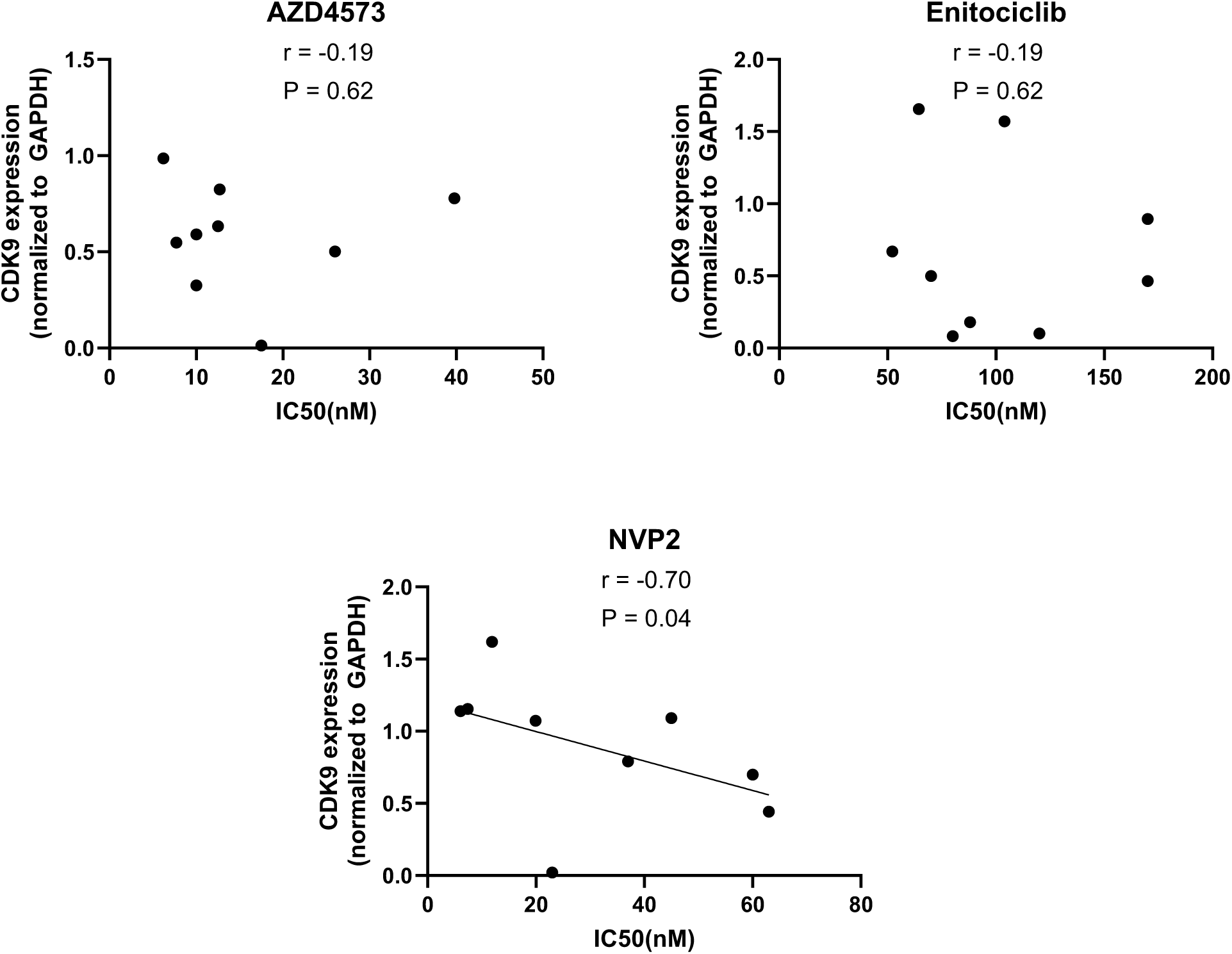
CDK9 protein expression (normalized to GAPDH loading control from concurrent protein lysates) plotted against the IC50 values of matched CRC cell lines or PDOs. Protein expression is estimated from immunoblot quantification. 9 models were included: HCT116, RKO, DLD-1, Lim1215, 22-006, 22-008, 23-007, 23-009, 23-010. All models were treated for 24 h with respective CDK9 inhibitors. Spearman correlations were performed and r values along with P values represented within the graphs. NVP2 linear regression was performed and line of best fit plotted. All analysis was performed with Graphpad Prism.

**Supplementary Figure S7.**
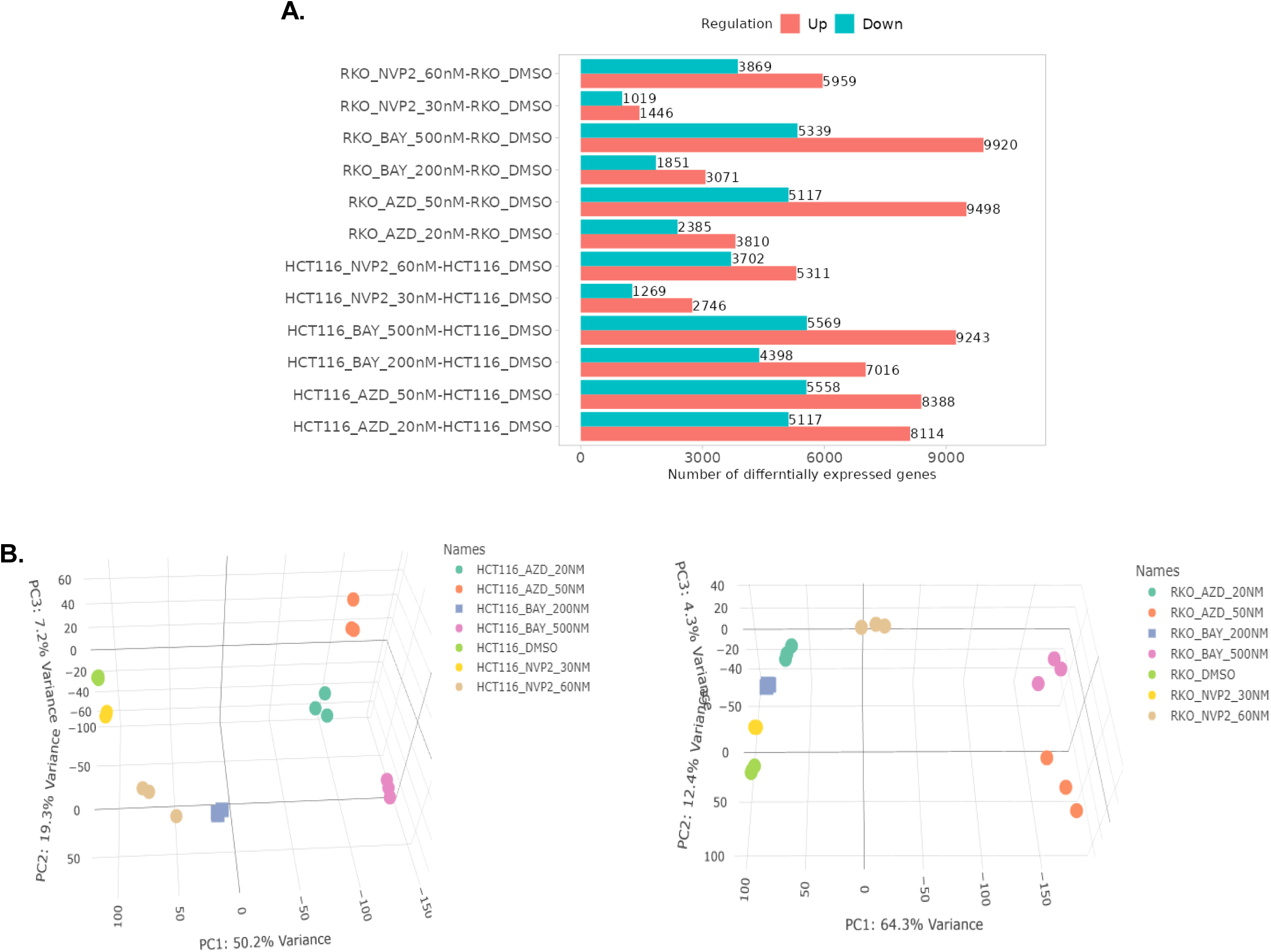
A) Total number of transcripts with significant increase or decrease in expression in CRC cells after treatment with CDK9 inhibitors for 24 h. Mean of 3 RNAseq replicates are shown. B) 3-D principal component analysis on transcriptomic profiles of HCT116 (left) and RKO (right) CRC cells after treatment with CDK9 inhibitors. Individual dots represent individual replicates of RNAseq.

**Supplementary Figure S8.**
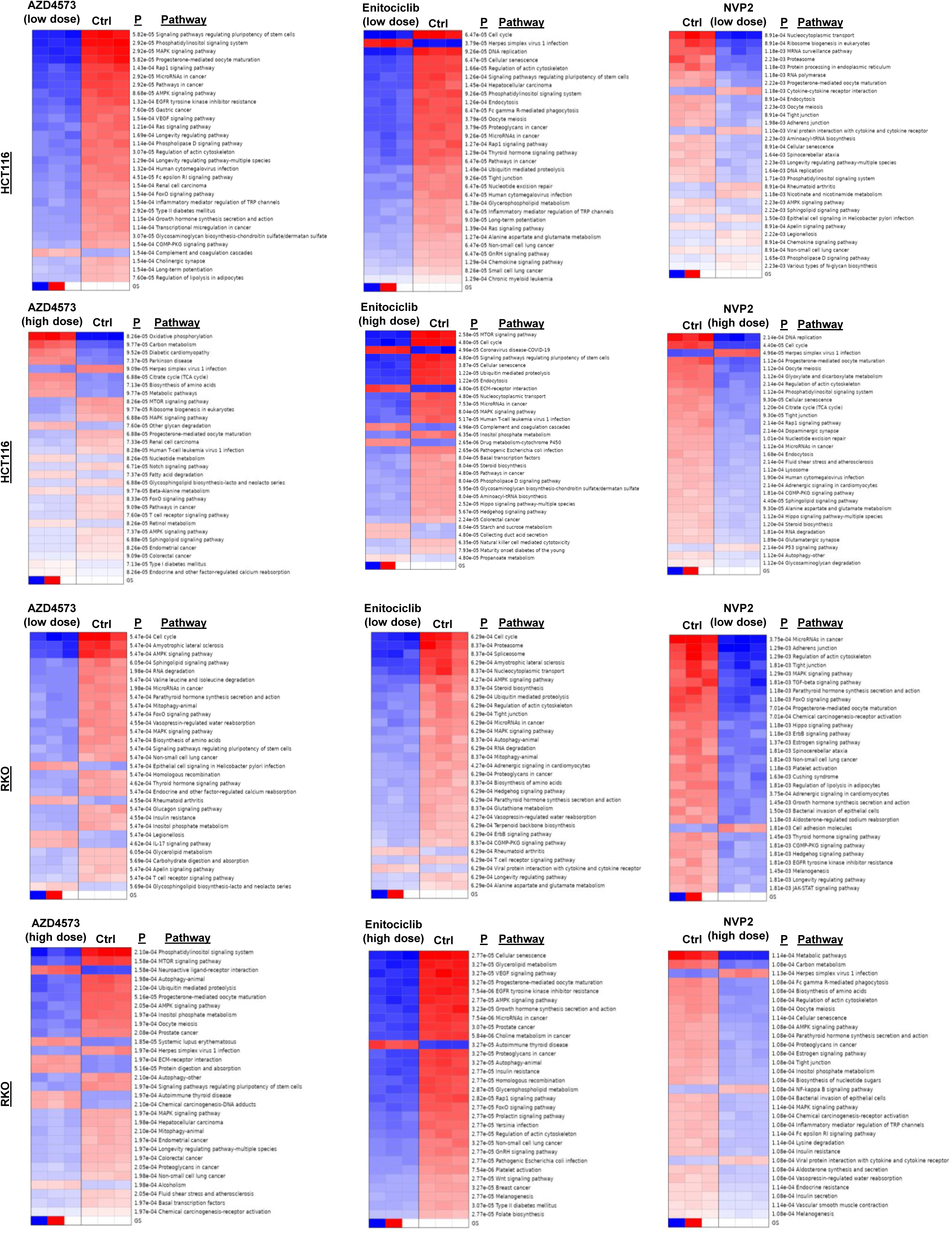
Parametric GSEA of indicated CRC cells treated with AZD4573 (20 or 50 nM), enitociclib (200 or 500 nM), or NVP2 (30 or 60 nM) for 24 h, as compared to DMSO control treatment. Three replicates of RNAseq are shown for each cell lines+treatment condition. Red = high, blue = low.

**Supplementary Figure S9.**
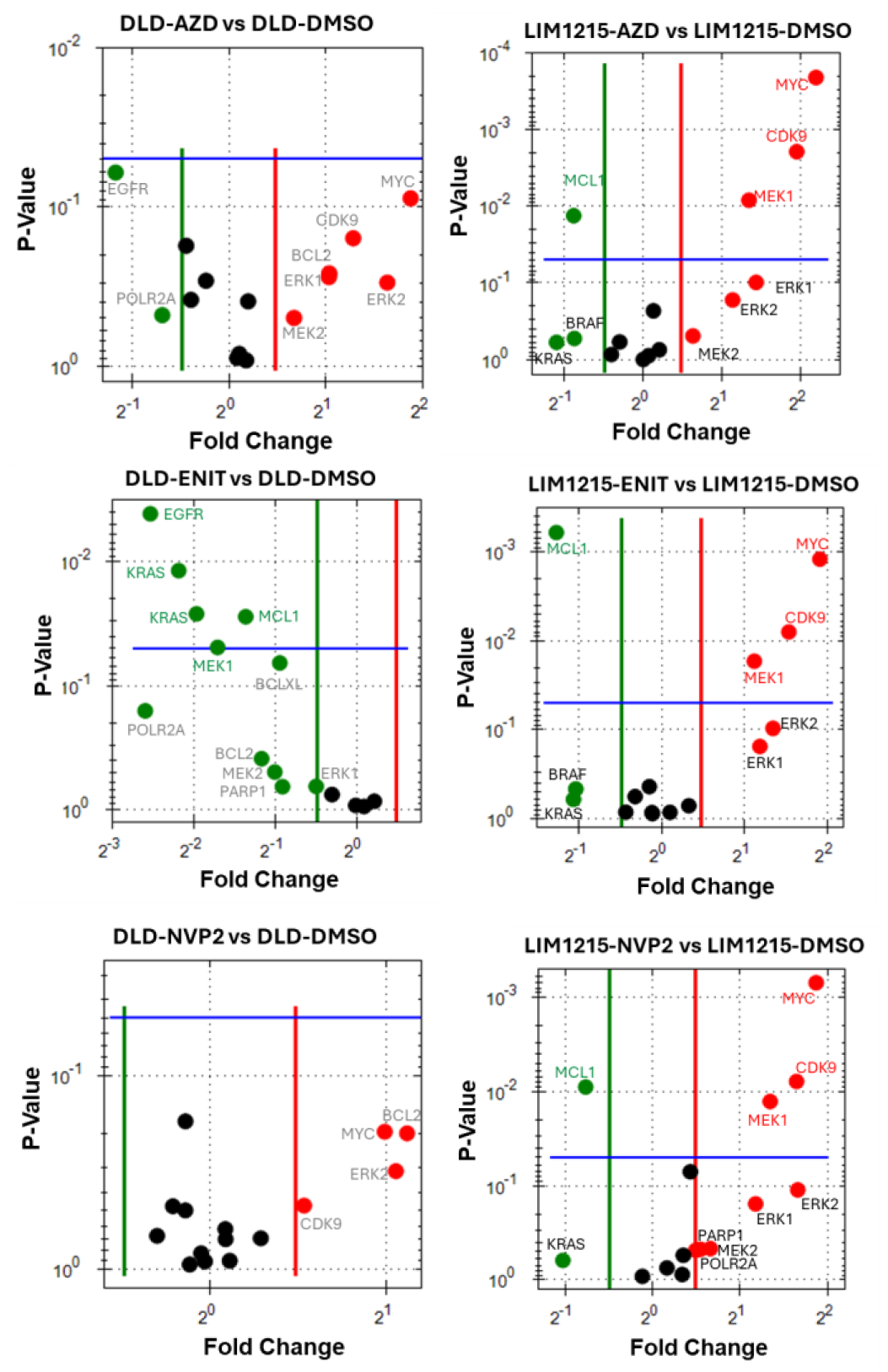
RT-qPCR expression levels of transcription genes and apoptosis genes in CRC cells treated with CDK9 inhibitors for 24 h, as compared to DMSO control treatment.

**Supplementary Figure S10.**
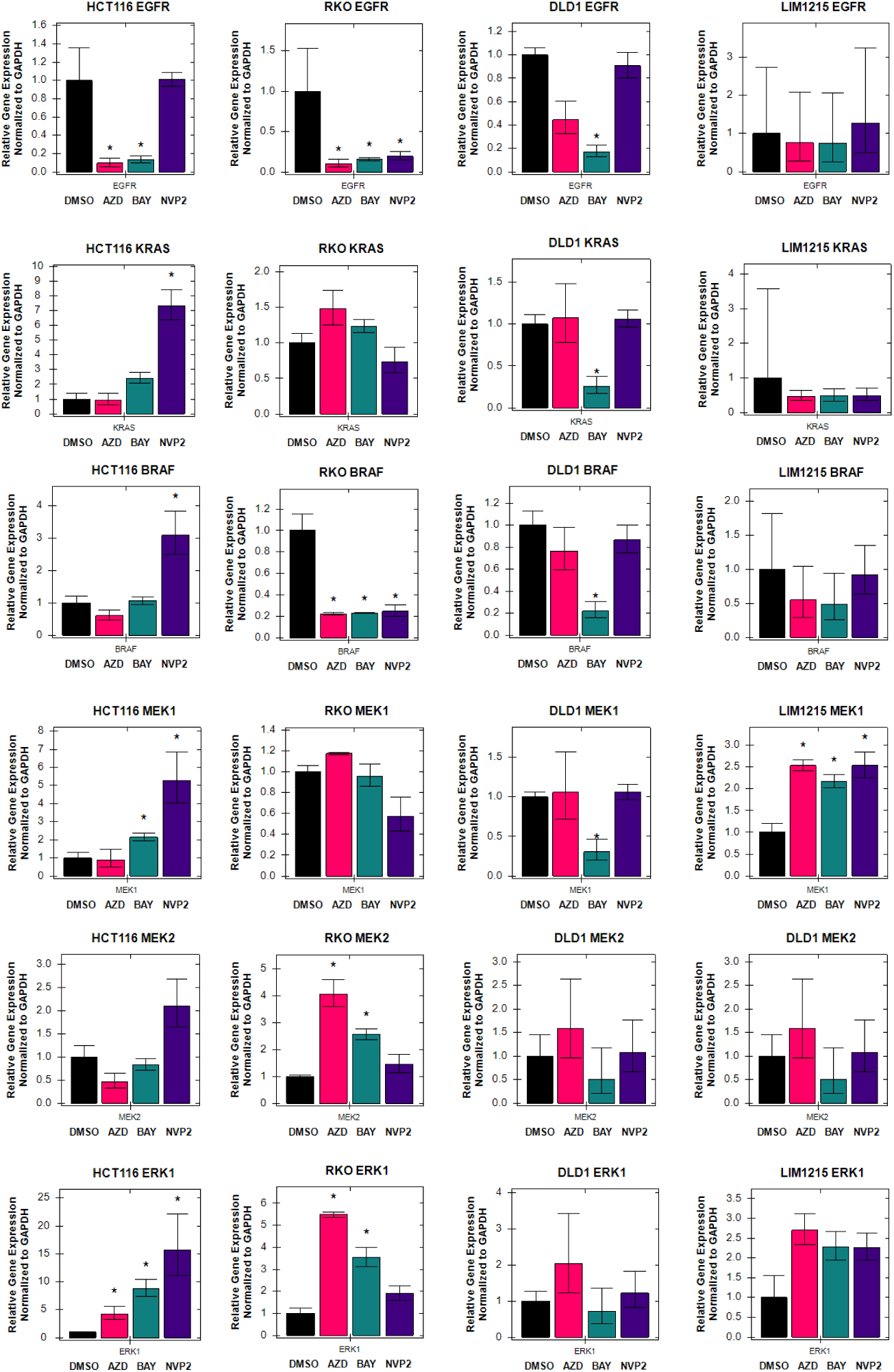
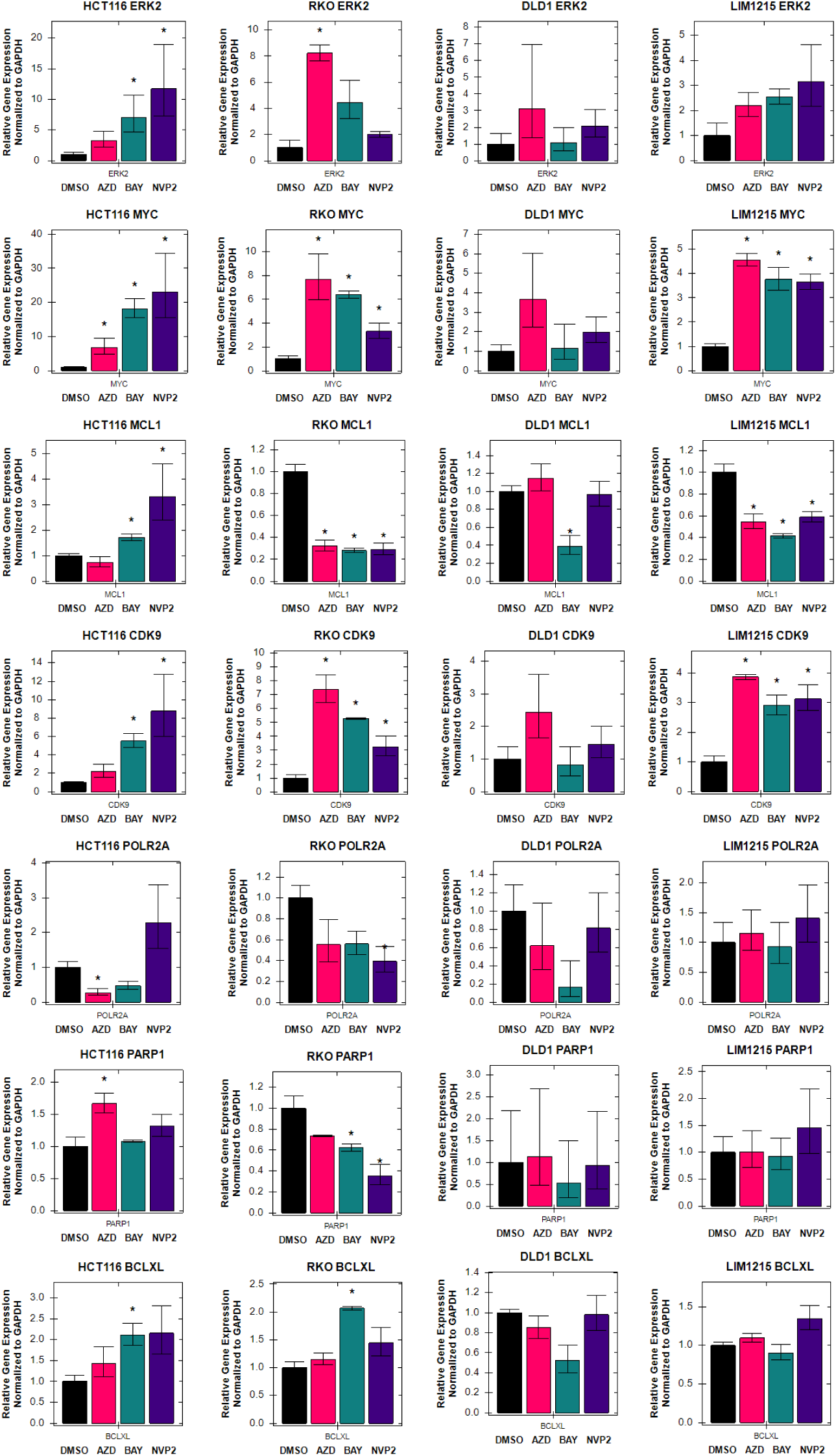
RT-qPCR expression levels of indicated transcripts in CRC cells treated with CDK9 inhibitors for 24 h, as compared to DMSO control treatment. RT-qPCR expression levels of indicated transcripts in CRC cells treated with CDK9 inhibitors for 24 h, as compared to DMSO control treatment.

**Supplementary Figure S11.**
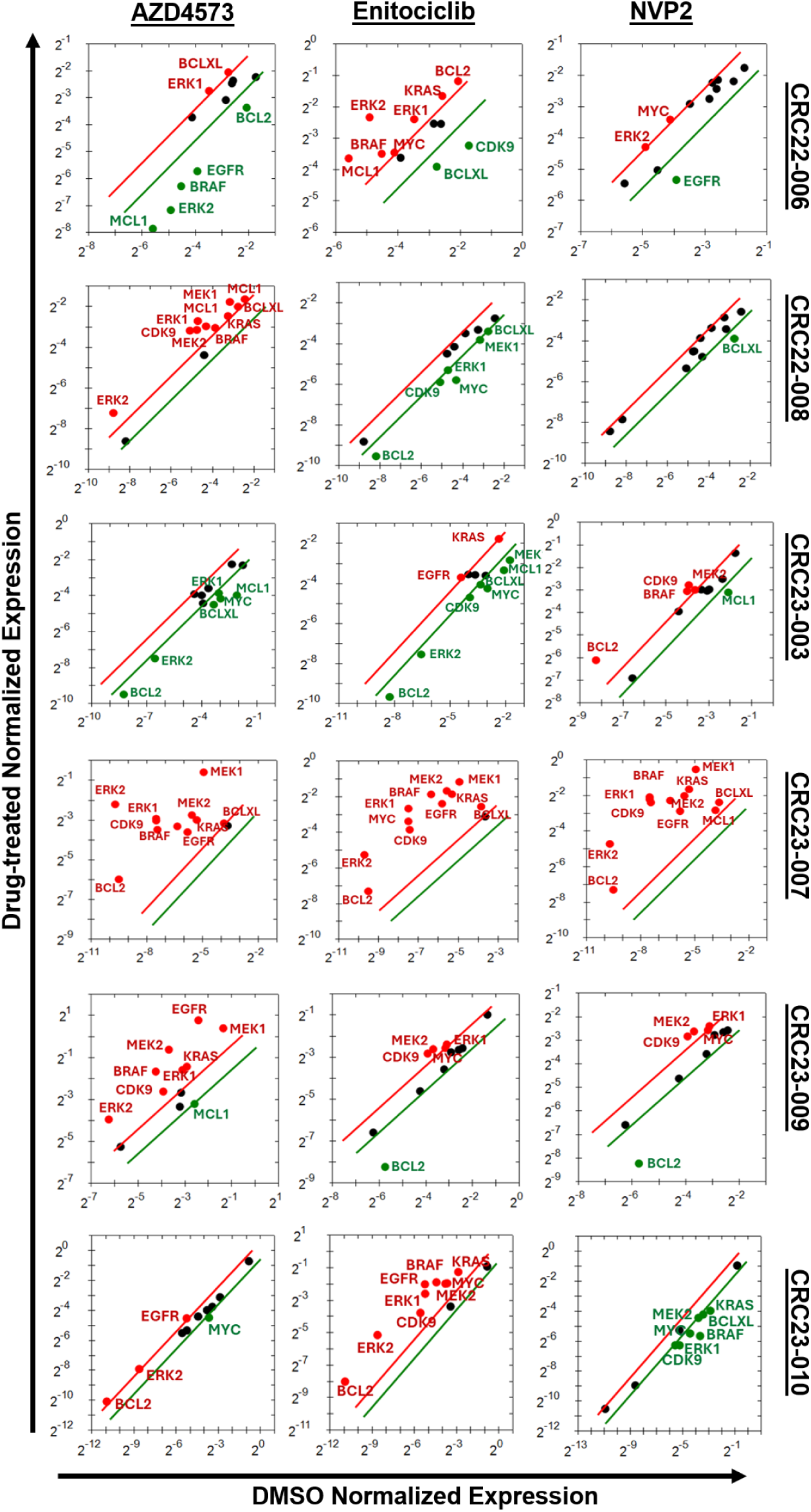
Relative expression level of key gene transcripts in PDOs after treatment with CDK9 inhibitors for 24 hours, as measured by RT-qPCR and normalized to GAPDH gene expression. PDOs were treated with either AZD4573 (50 nM), enitociclib (500 nM), or NVP2 (60 nM).

**Supplementary Figure S12.**
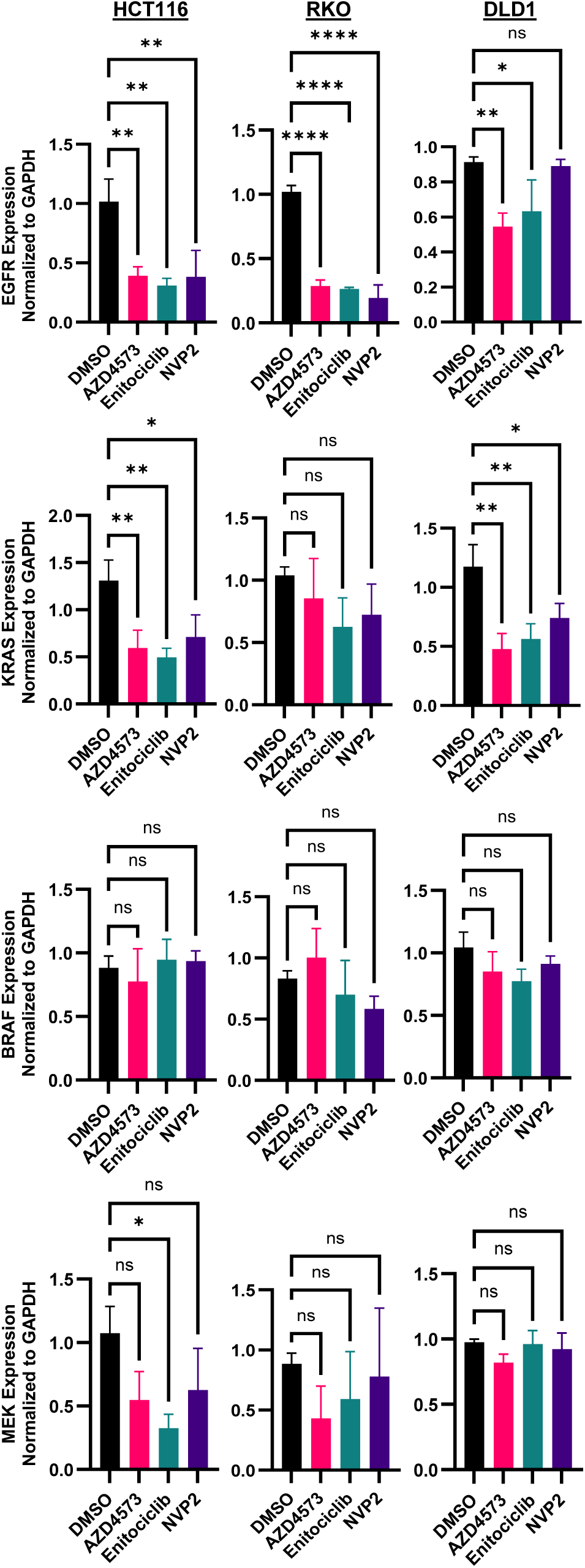
Quantification of immunoblots of MAPK pathway proteins, measured by densitometry, of indicated CRC cells after 24 h treatment with CDK9i’s. N=3 biological replicates. *: p<0.05, **: p<0.005.

**Supplementary Figure S13.**
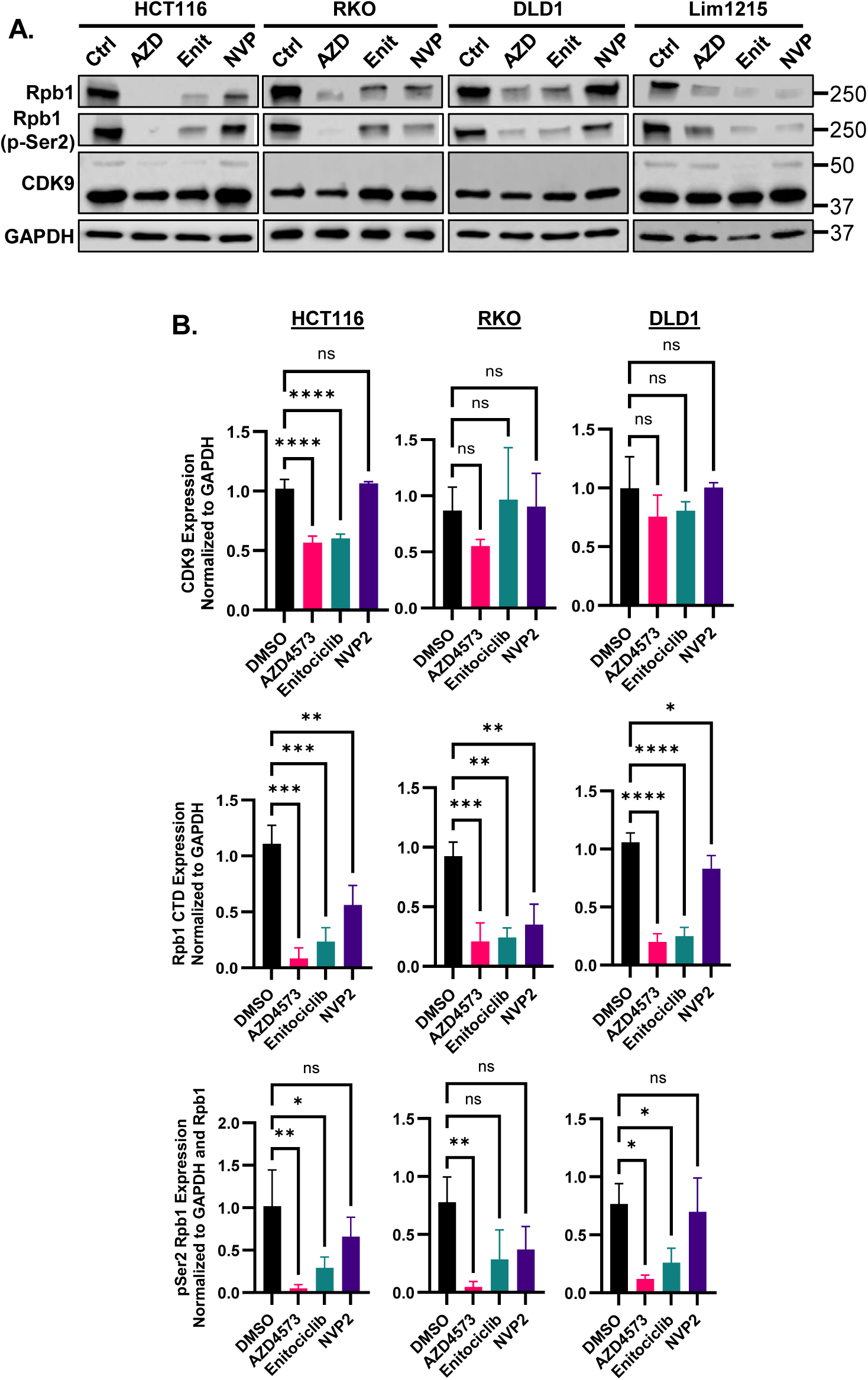
A) Representative immunoblots of Pol II and CKD9 proteins in CRC cells after 24 h CDK9 inhibitor treatment. B) Quantification of immunoblots of Pol II and CDK9 proteins, measured by densitometry, of indicated CRC cells after 24 h treatment with CDK9 inhibitors. N=3 biological replicates. *: p<0.05, **: p<0.005, ***: p<0.0005, ****: p<0.00005.

**Supplementary Figure S14.**
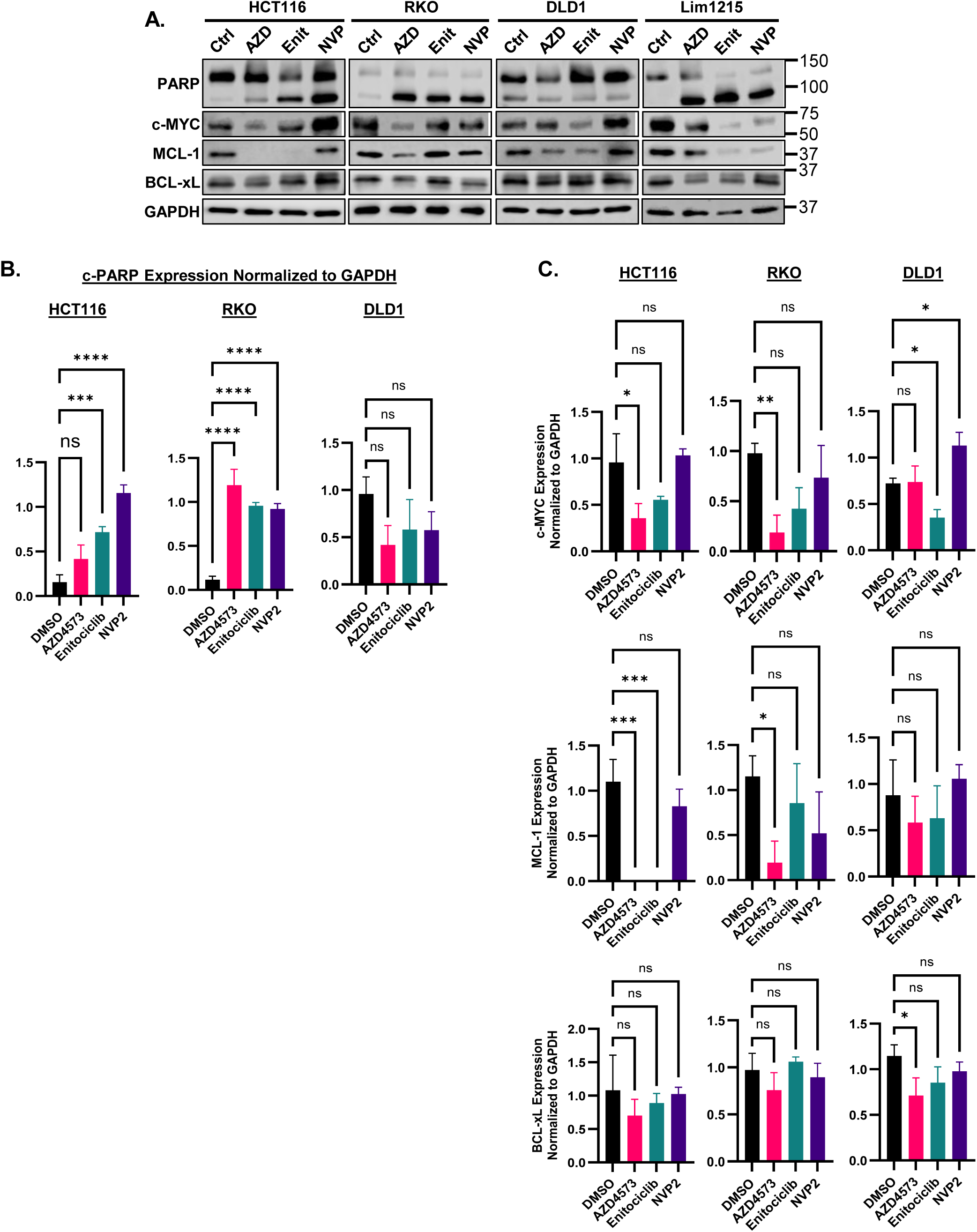
A) Representative immunoblots of PARP, c-MYC and Bcl-2 family proteins in CRC cells after 24 h CDK9 inhibitor treatment. B) Apoptosis as measured by densitometry of immunoblot bands of cleaved PARP in CRC cells treated with CDK9 inhibitors for 24 h. C) Quantification of immunoblots of c-MYC and Bcl-2 family proteins, measured by densitometry, of indicated CRC cells after 24 h treatment with CDK9 inhibitors. N=3 biological replicates. *: p<0.05, **: p<0.005, ***: p<0.0005, ****: p<0.00005.

**Supplementary Figure S15.**
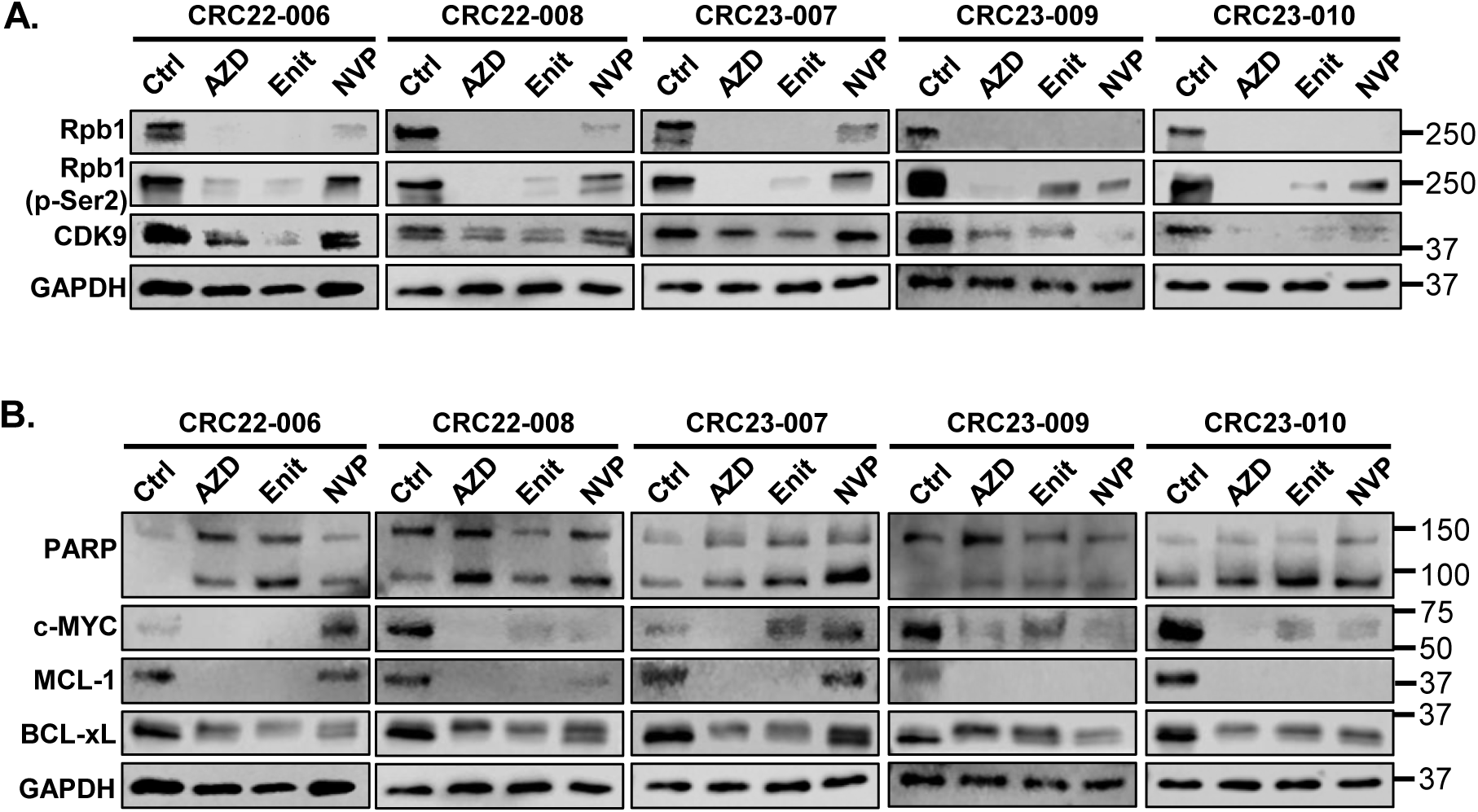
A) Representative immunoblots of Pol II and CKD9 proteins in CRC PDOs after 24 h CDK9 inhibitor treatment. B) Representative immunoblots of c-MYC and Bcl-2 family proteins in CRC PDOs after 24 h CDK9 inhibitor treatment.

**Supplementary Figure S16.**
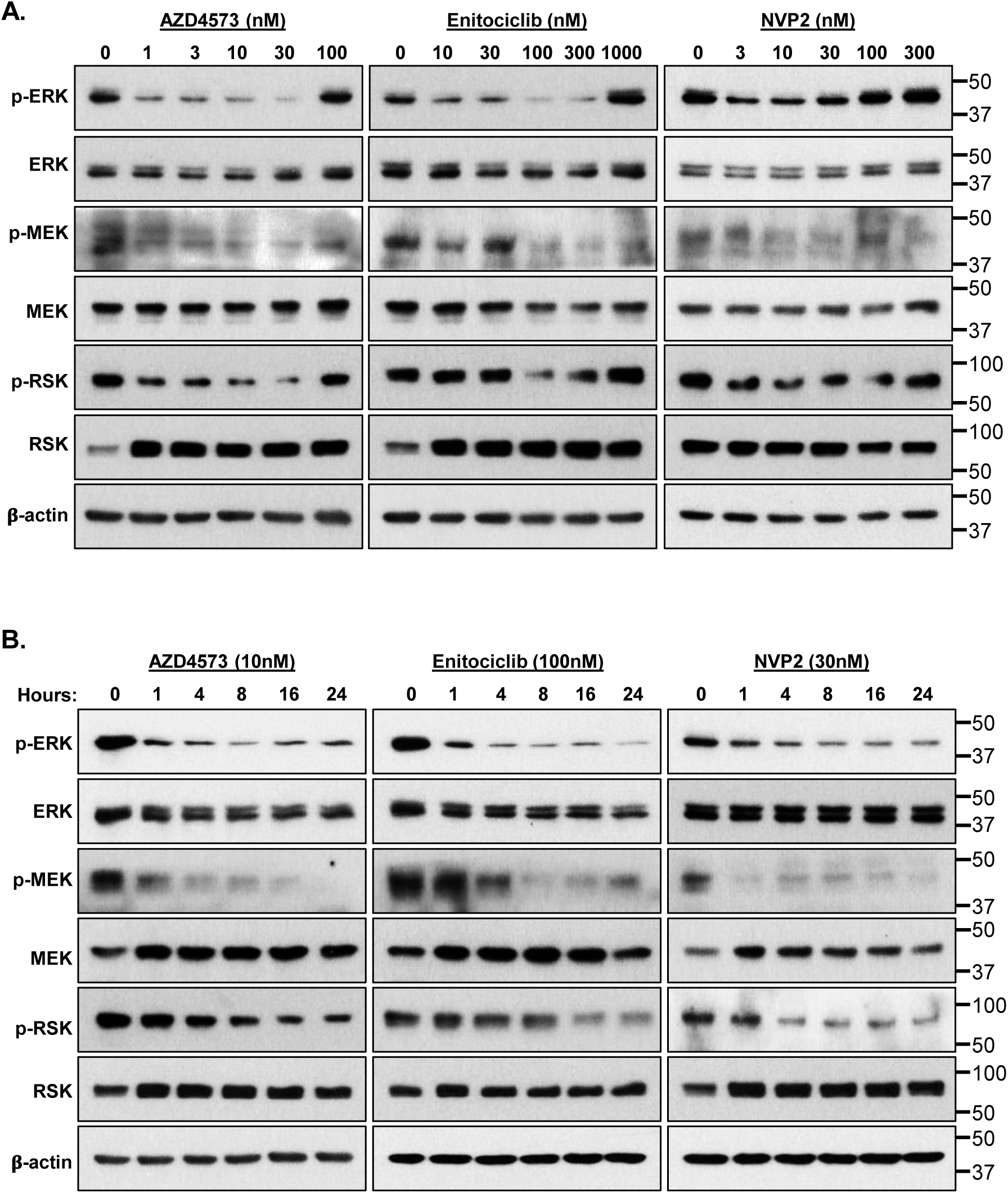
A) Representative immunoblots of MAPK signaling proteins and phospho-proteins in HCT116 cells after 24 h treatment with indicated concentrations of CDK9 inhibitors. B) Representative immunoblots of MAPK proteins in HCT116 cells treated with indicated CDK9 inhibitors over a 24-hour time course.

**Supplementary Figure S17.**
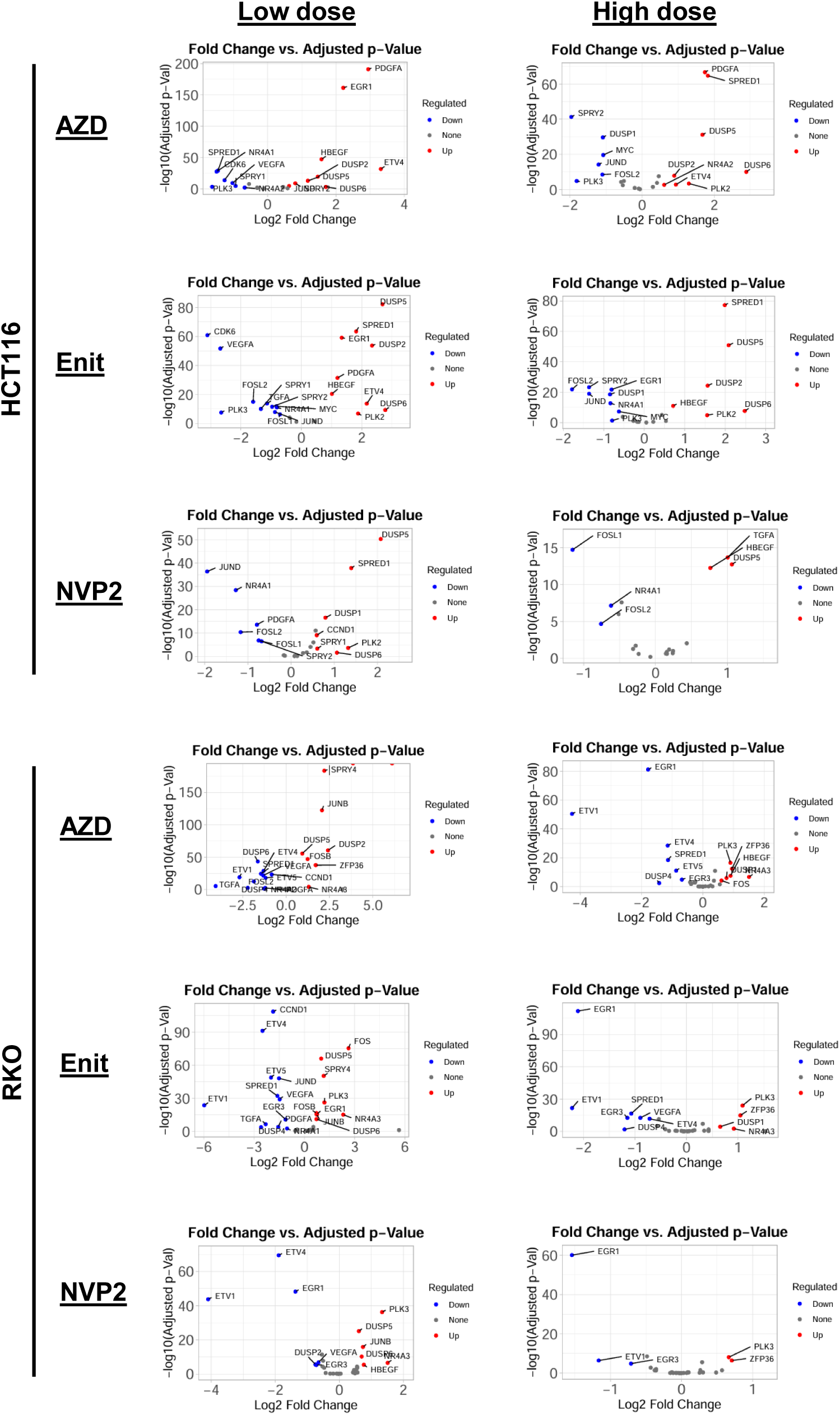
Gene expression changes of MAPK-regulated genes, after 24 h of CDK9 inhibitor treatment. Indicated cells were treated for 24 h with CDK9 inhibitor at either a low dose (AZD4573 20 nM, enitociclib 200 nM, NVP-2 30 nM) or high dose (AZD4573 50 nM, enitociclib 500 nM, NVP-2 60 nM).

