## Supplementary Tables 1 and 2 for "Cyclin-dependent kinase 9 inhibitors as oncogene signaling modulators in combination with targeted therapy for the treatment of colorectal cancer"

**Supplementary Table 1**

| **Source** | **Gene Symbol** | **Forward Primer** | **Reverse Primer** |
| --- | --- | --- | --- |
| IDT | ACTB | ATATGAGATGCGTTGTTA | AAGTATTAAGGCGAAGAT |
| IDT | BCL2 | GAGTGCTGAAGATTGATG | TCCTCTGTGATGTTGTATT |
| IDT | BCL2L1 | AAGCGTAGACAAGGAGAT | TAGGTGGTCATTCAGGTAA |
| IDT | BRAF | AGCAGTTACAAGCCTTCAA | GATATGGAGATGGTGATACAAG |
| IDT | CDK9 | CACTGGAGGTCTTGACTT | ATGAGATGCGTTCTGGAA |
| IDT | EGFR | CCAAGGCACGAGTAACAA | GGCAATGAGGACATAACCA |
| IDT | GAPDH | CTCTGGTAAAGTGGATATTGT | GGTGGAATCATATTGGAACA |
| IDT | KRAS | TCTAAGTGCCAGTATTCC | CACACCAACATTCACAAT |
| IDT | MCL1 | CCAGGCAAGTCATAGAAT | GAGGCTTACAGTCATAGTT |
| IDT | MAPK1 | TTGGATGTGGTGTTATGGAA | AAGCAGAGACGCAGAATG |
| IDT | MAPK3 | AGAATGTCATCGGCATCC | TACAGGTCAGTCTCCATCA |
| IDT | MAP2K1 | ATAGTCAACGAGCCTCCT | CCACTTCCTCAGCATCAG |
| IDT | MAP2K2 | GAACTCCTGGACTATATTGTGA | GCTTGATGAAGGTGTGGTT |
| IDT | MYC | CTCAAGTCATAACAATGCTAA | AATCAACAGTATCTCCTTCA |
| IDT | PARP1 | ACCACTTCTCCTGCTTCT | CTTCTCTGCCTTGCTACC |
| IDT | POLR2A | TAACAACTGGCTCCTCAT | TCCTGGTAAGTCTTAGAATCA |
| IDT | SOS1 | GCATTGTGGATGGAGGATA | TGAGAAGAAGGCAAGGATG |

**Supplementary Table 2**

| **Source** | **Item Name** | **Catalogue Number** | **Dilution** | **Incubation and time** | **Antigen Retrieval and time (Only IHC)** |
| --- | --- | --- | --- | --- | --- |
| Cell Signaling | Rpb1 CTD (4H8) | 2629 | 1:1000 | 4 C Overnight |  |
| Abcam | Anti-RNA polymerase II CTD repeat YSPTSPS (phospho S2) | AB5095 | 1:1000 | 4 C Overnight |  |
| Cell Signaling | CDK9 (C12F7) | 2316 | 1:1000 | 4 C Overnight |  |
| Cell Signaling | GAPDH (D16H11) XP | 5174 | 1:1000 | 4 C Overnight |  |
| Cell Signaling | PARP | 9542 | 1:1000 | 4 C Overnight |  |
| Cell Signaling | c-Myc (D84C12) | 5605 | 1:1000 | 4 C Overnight |  |
| Cell Signaling | Mcl-1 (D5V5L) | 39224 | 1:1000 | 4 C Overnight |  |
| Cell Signaling | Bcl-xL | 2764 | 1:1000 | 4 C Overnight |  |
| Cell Signaling | EGF Receptor (D38B1) XP | 4267 | 1:1000 | 4 C Overnight |  |
| Cell Signaling | K-Ras (E2M9G) | 71835 | 1:1000 | 4 C Overnight |  |
| Cell Signaling | B-Raf (D9T6S) | 14814 | 1:1000 | 4 C Overnight |  |
| Cell Signaling | MEK1/2 | 9122 | 1:1000 | 4 C Overnight |  |
| Cell Signaling | p44/42 MAPK (Erk1/2) | 9102 | 1:1000 | 4 C Overnight |  |
| Cell Signaling | Phospho-p44/42 MAPK (Erk1/2) (Thr202/Tyr204) | 9101 | 1:1000 | 4 C Overnight |  |
| Santa Cruz | Beta Actin | SC69879 | 1:2000 | 4 C Overnight |  |
| Invitrogen | Stabilized peroxidase conjugated goat Anti-mouse (H+L) | 324030 | 1:5000 | RT 1 hour |  |
| Biolegend | HRP Donkey Anti-Rabbit IgG | 406401 | 1:5000 | RT 1 hour |  |
| Abcam | Anti-Cleaved PARP1 antibody [E51] | AB32064 | 1:200 | 4 C Overnight | Citrate pH 6; 20 minutes |
| Abcam | ERK1 (phospho T202 + Y204) + ERK2 (phospho T185 + Y187) | AB223500 | 1:200 | 4 C Overnight | Citrate pH 6; 20 minutes |
| Abcam | Rabbit monoclonal [EPR3119Y] to Cdk9 | AB76320 | 1:200 | 4 C Overnight | Citrate pH 6; 20 minutes |
| Abcam | Rabbit polyclonal, Ki67 | AB15580 | 1:200 | 4 C Overnight | Citrate pH 6; 20 minutes |
| Vector Laboratories | Anti Rabbit IgG (H+L) biotinylated | BA1000 | 1:500 | RT 1 hour |  |
