## Supplementary Methods for "Cyclin-dependent kinase 9 inhibitors as oncogene signaling modulators in combination with targeted therapy for the treatment of colorectal cancer"

**Table of Contents**

| **Method** | **Page** |
| --- | --- |
| Organoid Establishment^1,2^ | 2 |
| Sub-culturing (Passaging) Organoids | 4 |
| Thawing Organoids | 6 |
| Freezing Organoids | 7 |
| WRN Conditioned Media Preparation | 8 |
| Organoid Complete Growth Media Formula | 9 |
| Transcriptomic Analysis of Cell Lines | 10 |
| Statistical Analysis of Gene Expression | 11 |
| RNAseq of Tumors and PDOs | 12 |
| RNAseq Data Analysis | 13 |
| WES of Tumors and PDOs | 14 |
| OneSeq-Based Sequencing of Tumors and PDOs | 15 |
| Targeted Sequencing of Tumors and PDOs | 16 |
| Reverse Transcriptase Quantitative PCR | 17 |

All organoid methods in this supplement were adapted from methods used in the following studies:

1.	Song, X., Shen, L., Tong, J., Kuang, C., Zeng, S., Schoen, R. E., *et al.* Mcl-1 inhibition overcomes intrinsic and acquired regorafenib resistance in colorectal cancer. *Theranostics* **10**, 8098–8110 (2020).

2.	Liu, C., Banister, C. E., Weige, C. C., Altomare, D., Richardson, J. H., Contreras, C. M., *et al.* PRDM1 silences stem cell-related genes and inhibits proliferation of human colon tumor organoids. *Proc Natl Acad Sci U S A* **115**, E5066–E5075 (2018).

**Human Colon Tumor Organoids Development from Tumor Tissue**

Prep (15 minutes):

a) Prepare 2 x 50 mL tubes of DPBS supplemented with 1% Pen/Strep and 100 ug/mL primocin, chill at 4 deg/on ice; prepare 50 mL of advanced DMEM/F-12 + 15 mM HEPES + 1% BSA (add 2 mL of a 25% BSA stock solution in water), this is for pre-wetting pipets to prevent organoids from sticking to them.
b) Thaw Matrigel in 4 deg; thaw an aliquot of organoid media in 4 ºC (not 37 ºC) overnight, add supplements once thawed (see “Organoids Complete Growth Media Formula”); thaw an aliquot of heat inactivated-FBS; Thaw 1 mL of gentle collagenase/hyaluronidase (Stem Cell Technologies). Thaw 1000🞨Y-27632, 500🞨Primocin.
c) Pre-warm 24-well plate. Pre-chill a box of sterile 200 uL pipet tips.
d) Gather 50-mL and 15-mL centrifuge tubes, 100-mm Petri dish, sterile sharp scissors or sterile razor blades, sterile tweezer/forceps.

Isolation and establishment (4 hours):

1. Harvest the surgically resected specimens or biopsy samples from hospital, at this point will be in 50 mL tube with basal media (DMEM+ Pen/Strep +ROCK inhibitor+primocin) on wet ice. Ideally use ~100 µL to 500 µL of tumor (range from a 5 mm cube to half of a 1 cm cube). Remove the tumor pieces that you will save for other experiments with sterile forceps.
2. Remove the media from tube, take care not to aspirate the tissue. Wash the tissue sample 5 X with ~10 mL cold DPBS+Pen/Strep+primocin. Each time, decant or aspirate the prior wash, add the new wash, swirl, and let sit on wet ice for 5 minutes. It is critical to wash carefully at this point to remove as much bacterial/fungal contamination as possible.
3. Transfer tissue pieces and small amount of DPBS to a non-adherent sterile petri dish, by pouring or with forceps. Mince the tissue to ~1 mm pieces. The pieces must be small enough to fit through a 10 mL serological pipet. I use a sterile razor blade to do this.
4. Wet the inside of a serological pipet with DMEM+HEPES+BSA and pipet transfer the minced tissue to a new 50 mL tube. Use DPBS as needed to wash/carry the extra small fragments into the new tube.
5. Dilute 1 mL of Gentle Collagenase/Hyaluronidase (Cat#: 07919, Stemcell Technology,) to 10 mL by adding 9 mL of DPBS+ Pen/Strep +primocin. Add 10µl of the stock ROCK inhibitor (RI, Y-27632, which is at 10mM stock) for a final ROCK Inhibitor concentration of 10uM. Add Primocin to a final concentration of 100µg/mL.
6. Add the freshly prepared 10 mL of Collagenase/Hyaluronidase+RI+Primocin to the tissue, incubate at 37C for 30 minutes with constant shaking. Alternately or in addition, manually shake the mixture every 10 minutes.
7. After incubation, pipette up and down 10 times with a pre-wetted (DMEM+HEPES+BSA) 10 mL serological pipette to break up the pieces and liberate tumor cell clusters.
8. Add 500µl of heat-inactivated FBS and let mixture sit for 2 minutes at room temperature. The big pieces will settle down to the bottom of the tube. You want to recover and process the SUPERNATANT.
9. Using a pre-wetted serological pipette (DMEM+HEPES+BSA), transfer the supernatant to a new 15 mL tube. The supernatant contains tumor single cells and tumor cell clusters. Keep the supernatant.
10. Centrifuge the cell suspension, 400x g, for 5 minutes to pellet the cells that you want.
11. (First Washing) Re-suspend / wash the cell pellet with 10mL of DBPS+ Pen/Strep +primocin. Pellet again as above.
12. Re-suspend / wash the cell pellet with 1mL of DPBS+ Pen/Strep +primocin. Remove a 20µL aliquot and put it on a petri dish or six well plate and count approximate how many crypts you have. You can use this count to estimate how many tumor clusters you will plate per gel droplet. (Second Washing) Add another 9mL of DPBS+ Pen/Strep +primocin to wash the tumor cells a second time. Centrifuge, aspirate the supernatant wash, and keep the tumor cell pellet.
13. Put the tumor cell pellet on ice to chill.
14. Re-suspend the cold tumor cell pellet in ice cold Matrigel. You want to achieve at least 50-200 clusters per 50µL of Matrigel. Keep everything on ice until you make the organoid drops.
15. Quickly remove the warm 24-well plate from the 37 °C CO_2_ incubator, and carefully dispense 50µL of Matrigel+organoid suspension/drop into the center of each well of a 24-well plate using pre-chilled 200-μL pipette tips.
16. Put it back in a 37 °C CO_2_ incubator for 10 min to solidify the Matrigel.
17. During the solidifying period, prepare enough volume of human colon tumor organoids complete growth media supplemented with 10 μM Y-27632 and 100 μg/mL Primocin and pre-warm the media.
18. Take the plate out of the incubator and add 500uL of prepared complete media to each well and incubate the plate at 37 °C CO_2_ incubator.
19. Observe the organoids daily and refresh media with 500 μL of complete growth media every 3-4 d after plating.

**Sub-culturing (Passaging) Organoids**

Prep (10 minutes):

1. Prewarm 24-well plates at 37 °C in incubator, use enough wells to accommodate your final number of passaged organoid cultures.
2. Prewarm media for final cultures: organoid media (Wnt/Rspondin/Noggin supplemented by conditioned L-WRN media, see separate recipe), without ROCK inhibitor (Y-27632) and without Primocin. Will need 500 μL per well of 24 well plate.
3. Prewarm basal medium (usually advanced DMEM/F12).
4. Prewarm TrypLE Express
5. Aliquot ice cold Matrigel, will need 50 μL per well of 24-well plate, or 25 μL per well of 48 well plate. Unclear how much for 96 well plate. These can all be aliquoted far in advance and frozen until needed for use. Keep these on ice throughout this passaging process.
6. Pre-chill sterile, clipped P200 pipet tips in -20 freezer.
7. Find ROCK inhibitor and keep on ice.

Passage (2 hours):

1. Use a P1000 pipette and disrupt the Matrigel in each well with old media, using a scraping motion all over the well bottom. Transfer Matrigel fragments and media to 15 mL tube. OK to pool wells that are the same samples/tissue of origin/etc.
2. Pipette up and down in 15 mL tube to break up the Matrigel pieces. Remove bubbles from the top.
3. Centrifuge at 400g for 5 minutes at room temperature. Organoids will be at the bottom, then Matrigel above them, then media.
4. Aspirate most of the media off without disrupting the Matrigel layer too much. Use a P200 to carefully remove the remaining top Matrigel layer and avoid removing organoids. This step requires experience to refine: the organoid-free portion of Matrigel is more transparent and the lower organoid-rich layer is more translucent/opaque. Try to keep as much organoid as possible
5. Add 1-2mL of TrypLE to each 15 mL tube. Pipette up and down to disrupt the pellet.
6. Place the tube in a 37 °C water bath for 10-15 min. Invert tubes several times at 5 min into incubation.
7. Pipette with a P200 to further disrupt the organoid pellet.
8. Add 10 mL basal medium (advanced DMEM) to quench, invert to mix, and centrifuge it at 4 °C at 400g for 5 min.
9. At this point there is almost no Matrigel left and there should be a pellet of organoids at the bottom that appears very similar to a 2D cell culture pellet. Carefully aspirate the supernatant without disturbing this pellet. Use a P20 to remove trace media/Matrigel if needed. Place tubes on ice (you should have an ice bucket already with Matrigel and ROCKi in room temperature rack).
10. Add an appropriate volume of Matrigel to each tube. Volume will depend on the intended split ratio and number of former wells combined to make each tube. Keeping tube on ice, pipette up and down with P200 to resuspend the organoid pellet in the Matrigel.
11. Remove prewarmed 24-well (or whichever size your using) plates. Plate 50 μL of Matrigel/organoid mixture into the center of each well, the goal is to make a droplet at the bottom which does not touch the side walls. Matrigel should start to set once it is contacting the pre-warmed plate.
12. Once organoids are on plates, place them back in 37 °C for 10 min to solidify.
13. If this is first passage, add ROCK inhibitor to prewarmed organoid media, final concentration 10 μM (usually add 80 μL of 500X or 40 μL of 1000X inhibitor to 40 mL tube of media). For subsequent passages of stably growing organoids, you can use straight organoid media.
14. Add 500 μL organoid media per well to solid Matrigel/organoid drops.
15. Change media every 3-4 days. Observe for density and organoid size. Passage organoids when appropriate.

**Thawing Organoids**

Prep (10 minutes):

8. Prewarm 24-well plates at 37 °C in incubator, use enough wells to accommodate your final number of passaged organoid cultures.
9. Prewarm media for final cultures: organoid media (Wnt/Rspondin/Noggin supplemented by conditioned L-WRN media, see separate recipe), without ROCK inhibitor (Y-27632) and without Primocin. Will need 500 μL per well of 24 well plate.
10. Prewarm basal medium (usually advanced DMEM/F12).
11. Aliquot ice cold Matrigel, will need 50 μL per well of 24-well plate, or 25 μL per well of 48 well plate. These can all be aliquoted far in advance and frozen until needed for use. Keep these on ice throughout this passaging process.
12. Pre-chill sterile, clipped P200 pipet tips in -20 °C freezer.
13. Find ROCK inhibitor and keep on ice.

Thaw and plate (0.5 hour):

16. Quickly thaw vial(s) of frozen organoids in 37 °C water bath, must thaw quickly and completely (within 2 min).
17. Transfer the cells to a 15 mL conical tube containing 5 mL organoid media + ROCKi, centrifuge at 400g for 5 minutes. Organoids will be at the bottom.
18. Carefully remove supernatant, then resuspend organoid pellet with required volume of Matrigel. You can observe a small sample of organoid clusters under microscope first, to estimate the density and Matrigel volume needed (target 100-200 clusters/50 μL Matrigel). If in doubt, prepare only 2-4 wells max per vial of organoids thawed, to prevent them from being too sparse in the initial passage.
19. Keeping the organoids + Matrigel mixture on ice, using pre-chilled, clipped P200 tips, drop 50 μL of organoid + Matrigel into the middle of each well in the prewarmed 24-w plate.
20. Once organoids are on plates, place them back in 37 °C for 10 min to solidify.
21. If this is the first passage, add ROCK inhibitor to prewarmed organoid media, final conc 10 μM (usually 80 μL of 500X or 40 μL of 1000X inhibitor to 40 mL tube of media). For subsequent passages of stably growing organoids, you can use straight organoid media.
22. Add 500 μL organoid media per well to solid Matrigel/organoid drops.
23. Change media every 3 days (usually). Observe for density and organoid size, passage when appropriate.

**Freezing Organoids**

Prep (10 minutes):

1. Prepare organoid freezing media (70% complete organoid media + 20% FBS + 10% DMSO + Primocin + Y-27632 (ROCK inhibitor)), chill at 4 °C.
2. Prewarm TrypLE Express, basal medium (usually advanced DMEM/F12).
3. Find ROCK inhibitor and keep on ice.

Resuspend and freeze (1 hour):

1. Use a P1000 pipette and disrupt the Matrigel in each well with old media, using a scraping motion all over the well bottom. Transfer Matrigel fragments and media to 15 mL tube. OK to pool wells that are the same samples/tissue of origin/etc.
2. Pipette up and down with P1000 in 15 mL tube to break up the Matrigel pieces. Remove bubbles from the top.
3. Centrifuge at 400g for 5 min at room temperature. Organoids will be at the bottom, then Matrigel above them, then media.
4. Aspirate most of the media off without disrupting the Matrigel layer too much. Use a P200 to carefully remove the remaining top Matrigel layer and avoid removing organoids. This step requires experience to refine: the organoid-free portion of Matrigel is more transparent and the lower organoid-rich layer is more translucent/opaque. Try to keep as much organoid as possible
5. TrypLE should be supplemented with 10 μM ROCK inhibitor (Y-27632) immediately prior to use, at this point. Add 1mL of TrypLE + ROCKi to each 15 mL tube. Pipette up and down to disrupt the pellet.
6. Place the tube in a 37 °C water bath for 10 min. Invert tubes several times at 5 min into incubation.
7. Add 10 mL basal medium (advanced DMEM/F-12) to quench, invert to mix, and centrifuge it at 4 °C at 400g for 5 min.
8. At this point there is almost no Matrigel left and there should be a pellet of organoids at the bottom that appears very similar to a 2D cell culture pellet. Carefully aspirate the supernatant without disturbing this pellet. Use a P20 to remove trace media/Matrigel if needed. Place tubes on ice (should have an ice bucket already with Matrigel and possibly ROCKi in it).
9. Resuspend in chilled organoid freezing media. We would suggest splitting a 50 uL organoid-Matrigel drop into no more than 3 or 4 different 1mL cryovials.
10. Aliquot organoids into cryovials and freeze overnight using a slow-freezing container in a -80 °C freezer.
11. Transfer vials to a liquid nitrogen or -150 °C freezer the following day for long-term storage.

**WRN Conditioned Media (WRN-CM) Preparation Procedure**

1. Revive the L-WRN (ATCC CRL-3276) cells from -150 °C, add directly to flask with DMEM+10% FBS (ATCC DMEM #30-2002, no antibiotics) for passage 0, then next day switch to L-WRN complete growth media (DMEM+FBS+hygromycin B+G418) as per manufacturer instructions.
2. Split one T75 flask of confluent L-WRN cells in culture medium without G418 and hygromycin B into two or three T75 flasks.
3. Incubate the T75 flasks for 3 or 4 days or until the cells become over-confluent and there is scant visible cell aggregates detached.
4. Remove media and rinse flasks with 3 mL fresh media (no antibiotics) and discard rinse.
5. Add 15 mL fresh media (no antibiotics)/flask and incubate flasks for 24 hours.
6. First Batch CM Collection. Remove the medium to a centrifuge tube. Add new medium (15 mL) to each flask. Centrifuge the conditioned medium at 2000g for 5 minutes and decant supernatant into a large, sterilized bottle. Store the conditioned medium at 4 °C.
7. Repeat step 6 every 24 hours. By this method, you will collect 2nd, 3rd and 4th conditioned medium collections. Centrifuge and add to same collection bottle.
8. Collect 5th to 8th and 9th to 12th conditioned media, if desired.
9. Keep collections 1-4 and 5-8 and 9-12 (batches 1, 2, and 3 respectively) separate from each other. Each batch can be frozen in aliquots or mixed with basal medium (Advanced DMEM/F12, 50%) plus additional supplements (see Complete Growth Media Formula) and then aliquoted/frozen.
10. Be sure to track which batch is being used for each experiment, in case of inter-batch variability in the culturing outcomes.

**Organoids Complete Growth Media Formula (constituted with WRN-CM)**

| **Component** | **Stock** | **Vol to add (total is 500 mL)** | **Final concentration** |
| --- | --- | --- | --- |
| WRN-CM (DMEM based) | N/A | 250 mL | 50% |
| Advanced DMEM/F12 | N/A | 218 mL | N/A |
| 50🞨B27 | 50X | 10 mL | 1🞨 |
| 100🞨N2 (only component that Kuo lab does not use?) | 100X | 5 mL | 1🞨 |
| N-Acetylcysteine | 500 mM | 1 mL | 1 mM |
| [leu-15]-GastrinⅠ | 100 μM | 50 μL | 10 nM |
| GlutaMax | 100X | 5 mL | 1🞨 |
| 100🞨HEPES | 100X | 5 mL | 1🞨 |
| 100🞨Pen/Strep | 100X | 5 mL | 1🞨 |
| Make 40 mL aliquots, freeze in -80 °C.  For experiments, thaw a 40 mL aliquot slowly in 4 °C overnight, then add the following: | | | |
|  | Stock | Vol to add (total Vol is 40 mL) | Final Conc |
| Nicotinamide | 2 M | 200 μL | 10 mM |
| EGF | 50 ug/mL | 40 μL | 50 ng/ml |
| SB202190 | 30 mM | 13.4 μL | 10 μM |
| A83-01 | 5 mM | 4 μL | 0.5 μM |
| PGE2 | 10 μM | 40 μL | 10 nM |
| Total final volume is 500 mL.  Add the following immediately before using in culture: | | | |
| Y-27632 | 10 mM (500X) |  | 10 μM |
| Primocin | 50 mg/mL (500X) |  | 100 μg/ml |

**Transcriptomic Analysis of Cell Lines Treated with a CDK9 Inhibitor Using RNA Sequencing**

Total RNA was extracted from 30 samples derived from human colorectal cancer cell lines (HCT116 and RKO) using the Direct-zol RNA MiniPrep Kit (Zymo Research, Cat # R2071) following the manufacturer's protocol. RNA concentration and purity were evaluated using a NanoDrop 2000 spectrophotometer (Thermo Fisher Scientific), with samples meeting quality thresholds of a 260/280 ratio >2.0 and a 260/230 ratio >2.0 selected for further analysis. RNA integrity was assessed using the Agilent Bioanalyzer through capillary electrophoresis, and samples with an RNA Integrity Number (RIN) meeting the quality control criteria were subsequently processed for library preparation.

RNA sequencing (RNA-seq) was performed on the NovaSeq 6000 platform (PE150) using poly-A capture and non-directional library preparation. The RNA-seq workflow comprised three main steps: library preparation, sequencing, and data analysis. mRNA was enriched from total RNA using Oligo(dT) magnetic bead enrichment, followed by ribosomal RNA depletion to obtain purified mRNA. Low-input eukaryotic mRNA library preparation was conducted (non-directional) with random fragmentation using Fragmentation Buffer (New England Biolabs, NEB) as per the manufacturer’s instructions. First-strand cDNA synthesis was performed using fragmented mRNA as a template, with random hexamer primers and reverse transcriptase. Second-strand cDNA synthesis was carried out using a custom buffer (NEBNext® Ultra™ RNA Library Prep Kit for Illumina®), dNTPs, RNase H, and Escherichia coli DNA polymerase I, employing nick-translation. The resulting cDNA library underwent purification, terminal repair, A-tailing, adapter ligation, size selection, and PCR amplification. Sequencing adapters, including P5/P7 (PCR amplification primers and flow cell binding regions), index sequences (for library multiplexing), and Rd1/Rd2 SP (Read 1/Read 2 sequencing primer binding regions), were added. The sequencing process initiates at the Rd1/Rd2 SP regions. Library concentration was initially measured using a Qubit 2.0 Fluorometer (Life Technologies) and diluted to 1.5 ng/µL. The insert size was assessed on an Agilent 2100 Bioanalyzer, and library quantification was refined using quantitative PCR (qPCR) to achieve a final library concentration of 2 nM. Libraries passing quality control were processed further based on their effective concentration and the desired data output. Sequencing was performed using the synthesis-by-sequencing (SBS) method. During sequencing, four fluorescently labeled dNTPs, DNA polymerase, and linker primers were introduced to the flow cell. As complementary strands were extended, fluorescently labeled dNTPs incorporated into the strand emitted fluorescence signals, which were captured by the sequencer. These optical signals were converted into sequencing peaks using computational software, providing the nucleotide sequence information for each fragment.

Raw sequencing data were processed into sequenced reads using CASAVA base calling, with the output recorded in FASTQ format, containing both sequence information (reads) and associated quality scores. Since sequencing processes are prone to machine errors, the quality of the raw data was evaluated, including an analysis of the sequencing error rate distribution, which reflects overall data quality. To minimize downstream analytical artifacts, low-quality reads and adapter-contaminated reads were removed during a filtering step to generate clean reads. Reads were excluded if they met any of the following criteria: (1) contained more than 10% ambiguous nucleotides (N > 10%); or (2) had low-quality scores (Qphred ≤ 20) across a significant portion of the sequence. Quality control checks included filtering, error rate analysis, and GC content distribution assessment, resulting in a high-quality dataset of clean reads for subsequent analyses.

Clean reads were aligned to the reference genome (hg38) using HISAT2, a fast and sensitive alignment tool optimized for RNA-seq data. HISAT2 leverages a hierarchical index strategy, comprising a large set of small graph FM (GFM) indexes covering the entire genome. These local indexes, combined with advanced multiple alignment strategies, enable the efficient mapping of reads, including those spanning multiple exons. The alignment process involved mapping sequences in stages: (1) reads were aligned to single exons; (2) reads spanning two exons were aligned using split mapping; and (3) reads spanning three or more exons were aligned through segmentation. Aligned regions were categorized as exonic, intronic, or intergenic. The majority of reads mapped to exons, reflecting active transcription. Intronic reads were attributed to either pre-mRNA contamination or intron retention events arising from alternative splicing mechanisms.

**Statistical Analysis of Differentially Expressed Genes and Gene Set Enrichment Analysis**

Reads that passed quality control, including removal of adapter sequences and low-quality reads, and were successfully mapped to the reference genome, were used for gene expression evaluation and quantification. Gene expression correlation analyses, including principal component analysis (PCA) and hierarchical clustering, were performed to assess the consistency among biological replicates and evaluate their similarity across treatment groups and doses. Differential expression analysis between two or more sample groups was conducted using DESeq2 (https://www.bioconductor.org/packages). This method employs the Wald test with a negative binomial distribution to identify differentially expressed genes (DEGs). To account for multiple comparisons, the Benjamini-Hochberg method was applied to adjust p-values and control the false discovery rate (FDR). Gene Ontology (GO) analysis and gene set enrichment analysis (GSEA) of the DEGs were performed using Parametric Analysis of Gene Set Enrichment (PAGE) implemented in the PGSEA package (https://www.bioconductor.org/packages). The enrichment significance was determined using Fisher's exact test. These analyses provided insights into the functional categories and biological pathways associated with the identified DEGs.

**RNA Sequencing for Characterization of Tumors and Patient-Derived Organoids**

Total RNA was extracted from five patient primary tumors and six corresponding patient-derived organoids (passages 5–10, one replicate each). Five organoids were derived from the primary tumors (C226T, C228T, C233T, C237T, and C2310T), while one organoid (C239T) lacked a corresponding primary tumor sample due to limited tissue availability from colonoscopy resection. RNA extraction was performed using the Direct-zol RNA MiniPrep Kit (Zymo Research, cat # R2071) according to the manufacturer’s protocol. RNA concentration and purity were assessed using a NanoDrop spectrophotometer (ThermoFisher). Samples meeting quality thresholds of a 260/280 ratio >1.8 and a 260/230 ratio >1.0 were further analyzed for RNA integrity using an Agilent Bioanalyzer. RNA Integrity Numbers (RINs) were determined through capillary electrophoresis, and only samples passing quality control (adequate RIN values and distinct 28S and 18S rRNA peaks) with sufficient RNA concentration were processed for library preparation.

**RNA Sequencing and Data Analysis**

RNA sequencing (RNA-seq) was performed on the NovaSeq 6000 platform (PE150) using poly-A capture and non-directional library preparation, employing the same methodology as used for cell line RNA-seq. Library preparation and sequencing followed established protocols. Data analysis employed the same pipeline, with quality control metrics including base calling with CASAVA and filtering of low-quality reads (Qphred ≤ 20). Clean reads, which passed quality control and were successfully mapped to the reference genome (hg38) using HISAT2, were further processed for gene expression evaluation and quantification. Gene expression correlation analyses, including principal component analysis (PCA) and hierarchical clustering, were conducted to evaluate the relationship between biological replicates and assess the similarity between organoids and their respective parental tumors. The analyses also examined whether tumor and organoid samples formed distinct clusters, reflecting their in vivo (tumor) or in vitro (organoid) environments. Additional factors considered included the absence of the extracellular matrix and non-epithelial cellular components, such as immune and stromal cells, in organoids compared to the primary tumor samples. Spearman correlation analysis was performed to quantify the similarity in gene expression profiles between each patient tumor and its corresponding patient-derived organoid.

**Whole Exome Sequencing of Patient Tumors and Organoids**

Genomic DNA was extracted from five primary tumor samples and six corresponding patient-derived organoids (passages 5–10, one replicate each) using the Quick-DNA/RNA™ MiniPrep Plus Kit (Zymo Research) following the manufacturer’s protocol. Among the six organoids, five were derived from the primary tumors (C226T, C228T, C233T, C237T, and C2310T), while one organoid (C239T) lacked a corresponding tumor sample due to limited tissue availability from colonoscopy resection. DNA concentration and purity were assessed with a NanoDrop spectrophotometer (ThermoFisher), ensuring a 260/280 ratio >1.8 and a 260/230 ratio >1.0. Electrophoretic quality was evaluated using 1% agarose gel electrophoresis, and DNA samples passing quality control were used for library preparation.

Genomic DNA was randomly sheared into fragments of 180–280 bp, followed by end-repair, A-tailing, and ligation with Illumina adapters. Adapter-ligated fragments underwent PCR amplification, size selection, and purification. Hybridization capture was performed using biotin-labeled probes to target exonic regions, followed by streptavidin-coated magnetic bead enrichment. Non-hybridized fragments were washed out, and hybridized fragments were enriched through PCR. Library concentration and size distribution were evaluated using Qubit fluorometry, real-time PCR, and a bioanalyzer, after which libraries were pooled and sequenced on Illumina platforms using a paired-end 150 bp (PE150) strategy. Exome sequencing targets the coding regions of the genome, offering a cost-effective alternative to whole-genome sequencing while focusing on regions associated with most known disease-related variants. Raw sequencing data, stored as fluorescence image files, were processed into short reads using base calling. The resulting reads, in FASTQ format, contained sequence information and base quality scores. Data quality control was performed using fastp (v0.23.1) to remove low-quality reads (Phred quality score ≤20) and adapter contamination.

Clean reads were aligned to the human reference genome (hg19) using the Burrows-Wheeler Aligner (BWA, v0.7.17). Aligned reads were processed with Picard (v2.18.9) to mark duplicates and with Sambamba (v1.0.0) for sorting and computing sequencing statistics, such as coverage and depth. Somatic SNP, InDel, and CNV calling were performed using the Genome Analysis Toolkit (GATK, v4.3.0), Mutect2, and Control-FREEC (v11.4). Annotation of variants was conducted with ANNOVAR, leveraging RefSeq and GENCODE databases to identify genomic regions, protein-coding changes, allele frequencies, and deleteriousness predictions. Statistical analyses were performed to assess somatic variants. This comprehensive workflow provided detailed insights into genomic variations, enabling the characterization of patient tumors and derived organoids.

**OneSeq-Based Sequencing for Tumor Characterization**

A complementary approach for tumor characterization involved sequencing with sex-matched reference DNA using Agilent OneSeq. For this method, either matched adjacent normal tissue or Agilent OneSeq Human Reference DNA (male: 5190-8848, female: 5190-8850) was used alongside tumor and organoid DNA. Library preparation was performed using the SureSelectXT platform (Agilent Technologies), and target enrichment was achieved through the OneSeq system. The OneSeq approach integrates ClearSeq catalog gene panels, SureSelect Exomes, Clinical Research Exome (CRE), Focused Exome, and the Inherited Disease Research Panel for targeted sequencing. It utilizes genome-wide baits to detect copy number variations (CNVs) and loss of heterozygosity (LOH), alongside user-defined baits for specific Agilent exomes, genes, or custom panels for identifying single nucleotide variants (SNVs) and indels. This method enables comprehensive sequencing and analysis of tumor and matched reference DNA, allowing for simultaneous genome-wide CNV, point mutation, and indel detection within a single assay. OneSeq provides a functional copy number resolution of 300 kb across the genome, with a higher resolution of 25–50 kb in clinically relevant regions, as defined by ClinGen. It also targets genomic regions with high minor allele frequency SNPs, enabling the detection of copy-neutral LOH at a resolution of 5 Mb. Data analysis was conducted using Agilent’s SureCall software, which facilitates the identification of CNVs, LOH, SNVs, and indels, providing a comprehensive genomic profile of the tumor samples. This approach enables precise and high-resolution characterization of patient tumors and patient-derived organoids.

**Targeted Sequencing of CRC Panel-Based Mutations**

To genotype colorectal cancer (CRC) patients and their derived organoids, the Ion Torrent AmpliSeq Colon Cancer Panel (Thermo Fisher Scientific) was employed as an alternative to whole-genome sequencing and the OneSeq method. This targeted sequencing approach was specifically utilized to validate critical CRC mutations. The Ion Torrent sequencing workflow was supported by an in-house analysis pipeline designed to streamline the next-generation sequencing (NGS) process by amplifying specific genomic regions of interest. Library preparation was performed using the Ion AmpliSeq Library 96LV Kit 2.0 (Life Technologies) in combination with the Colon Cancer Panel. Quality control of the prepared library was conducted using the Ion Sphere Quality Control Kit, following the manufacturer’s protocol. The emulsion PCR (emPCR) reaction targeted 10–30% of template-positive Ion Sphere particles (ISPs). Prior to loading onto 316 chips (100 Mb output), sequencing primers and polymerase were added to the final enriched ISPs.

The Ion AmpliSeq platform employs an ultrahigh multiplex polymerase chain reaction (PCR) strategy to selectively amplify targeted genomic regions, ensuring efficient and sensitive sequencing of CRC-specific gene panels and custom DNA hotspots. This panel covers key genes implicated in CRC, including AKT, ALK, AR, ARAF, BRAF, CDK4, CDKN2A, CHEK2, CTNNB1, EGFR, ERBB2, ERBB3, ERBB4, ESR1, FGFR, FLT3, GNAI1, GNAQ, GNAS, HRAS, IDH1, IDH2, KIT, KRAS, MAP2K1, MAP2K2, MET, MTOR, NRAS, NTRK1, NTRK2, NTRK3, PDGFRA, PIK3CA, PTEN, RAF1, RET, ROS1, SMO, and TP53. These genes were analyzed for single nucleotide variants (SNVs), while CNVs were assessed for a subset of genes, including ALK, AR, CD274, CDKN2A, EGFR, ERBB2, ERBB3, FGFR, KRAS, MET, PIK3CA, and PTEN. Sequencing was performed on the Ion PGM system (Life Technologies), and the resulting data were analyzed using Torrent Suite Software V3.2 (Life Technologies). The analysis included variant calling, annotation, and downstream interpretation, enabling the identification of significant mutations and actionable insights from the sequencing data. This targeted sequencing method provided a focused, efficient, and cost-effective approach for characterizing CRC-specific genetic alterations.

**Reverse Transcriptase Quantitative Real-Time PCR (RT-qPCR)**

Total RNA was isolated from patient tissue specimens, cell lines, and patient-derived organoids using the Direct-zol RNA MiniPrep RNA Isolation Kit (Zymo Research, Cat # R2071) according to the manufacturer’s protocol. RNA quantity and quality were assessed using a Nanodrop 2000 (Thermo Fisher), and RNA samples that met the following criteria were used for RT-qPCR analysis: a 260/280 ratio greater than 2.0 and a 260/230 ratio greater than 2.0. Two micrograms of total RNA were reverse-transcribed into complementary DNA (cDNA) using the iScript™ Advanced cDNA Synthesis Kit (Bio-Rad, cat # 1725037) according to the manufacturer’s instructions. The cDNA synthesis reaction was performed in a final volume of 20 µL in a T100 Thermal Cycler (Bio-Rad). The reaction included iScript Advanced Reverse Transcriptase, which consists of RNase H+ Moloney Murine Leukemia Virus (MMLV) reverse transcriptase and RNase inhibitor. The reaction was incubated at 42°C for 30 minutes, followed by reverse transcriptase inactivation at 95°C for 1 minute. The synthesized cDNA was stored at -20°C after being diluted 10-fold with nuclease-free water prior to qPCR analysis. For qPCR amplification, SsoAdvanced™ Universal SYBR Green Supermix (Bio-Rad, cat # 1725270) was used according to the manufacturer’s protocol. Reactions were carried out on a CFX Opus 384 Real-Time PCR System (Bio-Rad) with the following cycling conditions: initial denaturation at 95°C for 30 seconds, followed by 40 cycles of amplification (denaturation at 95°C for 15 seconds, primer annealing at 60°C for 30 seconds). A melt curve analysis was conducted from 65°C to 95°C with 0.5°C increments for 2-5 seconds per step. The primers used for each gene are listed in Supplementary Table S1.
